# Resolving a Systematic Error in STARR-seq for Quantitative Enhancer Activity Mapping

**DOI:** 10.1101/2020.10.20.346908

**Authors:** Longjian Niu, Jing Wan, Jialei Sun, Yingzhang Huang, Na He, Li Li, Chunhui Hou

## Abstract

STARR-seq assesses millions of fragments in parallel measuring enhancer activity quantitatively. Here we show that STARR-seq is critically flawed with a systematic error in the cells of *Arabidopsis thaliana (A. thaliana)*. Large amount of self-transcripts (STs) is lost during reverse transcription because these STs are polyadenylated after alternative polyadenylation sites (APAS) inside the test sequences. We solved this problem by using specially designed primer and recovered self-transcribed sequences independent from the PAS usage. In *A. thaliana*, we identified active enhancers and also enhancers quiescent in their endogenous genomic loci. Different from traditional STARR-seq identified enhancers, enhancers identified by new method are highly enriched in sequences proximal to the 5’ and 3’ ends of genes, and their epigenetic states correlate with gene expression levels. Our solution applies to methods based on self-transcript quantification. In addition, our results provide an invaluable functional enhancer activity map and insights into the functional complexity of enhancers in *A. thaliana*.

## Introduction

Enhancers are DNA sequences bound by transcription factors to increase gene transcription rate in a distance- and orientation-independent manner^1-3^. Enhancers are difficult to predict because of the lack of specific location relationship between enhancers and their target genes^4-7^. Chromatin marks including DNA accessibility, histone modification and transcription factor or cofactor binding have been used to characterize enhancers^8-17^. However, these methods do not provide the direct functional evidence or quantitative activity for enhancers.

STARR-seq is a functional test that measures enhancer activity in a direct, quantitative, and genome-wide manner^18^. This method has been adapted and widely used for enhancer analysis^19-24^. The accuracy of STARR-seq strictly relies on the full recovery of self-transcribed mRNAs initiated from the promoter of reporter gene. A systematic error may cause severe defect in the reliability of STARR-seq for both regulatory elements identification and activity quantification. In fact, previous work had revealed two major systematic errors in STARR-seq in human cells^19^.

In the original design of STARR-seq, it has been considered that transcription may be initiated inside the test DNA sequence instead of from the promoter of the reporter gene^18^. Self-transcripts (STs) initiated from the inside of a test DNA sequence is not expected to be recovered, and this loss does not affect the reliability of STARR-seq for enhancer identification and activity quantification^18^. However, if self-transcripts initiated from the promoter of reporter gene were not recovered, then the quantification of the enhancer activity of a test DNA sequence becomes problematic.

In the design of STARR-seq, mRNAs transcribed from the test DNA sequences are supposed to be polyadenylated after a polyadenylation site (PAS) placed at the end of the reporter gene in the plasmid construct. Since this PAS is designed to be used for the polyadenylation of self-transcripts, we refer to this PAS “designated PAS (DPAS)” (Fig. 1). However, one, even several, alternative PAS (APAS) may exist inside a test DNA sequence and the potential effects on the enhancer identification and activity quantification have not been considered when the original STARR-seq was designed. Theoretically, the usage of APAS for mRNA polyadenylation leads to the loss of these self-transcripts because the sequence that is complementary to reverse primer (RV) is lost (Fig. 1, right side in Box a). The loss of these self-transcripts may compromise the reliability of methods that are based on the quantification of STs which include both STARR-seq and other massively parallel reporter assays (MPRAs).

**Fig. 1.**
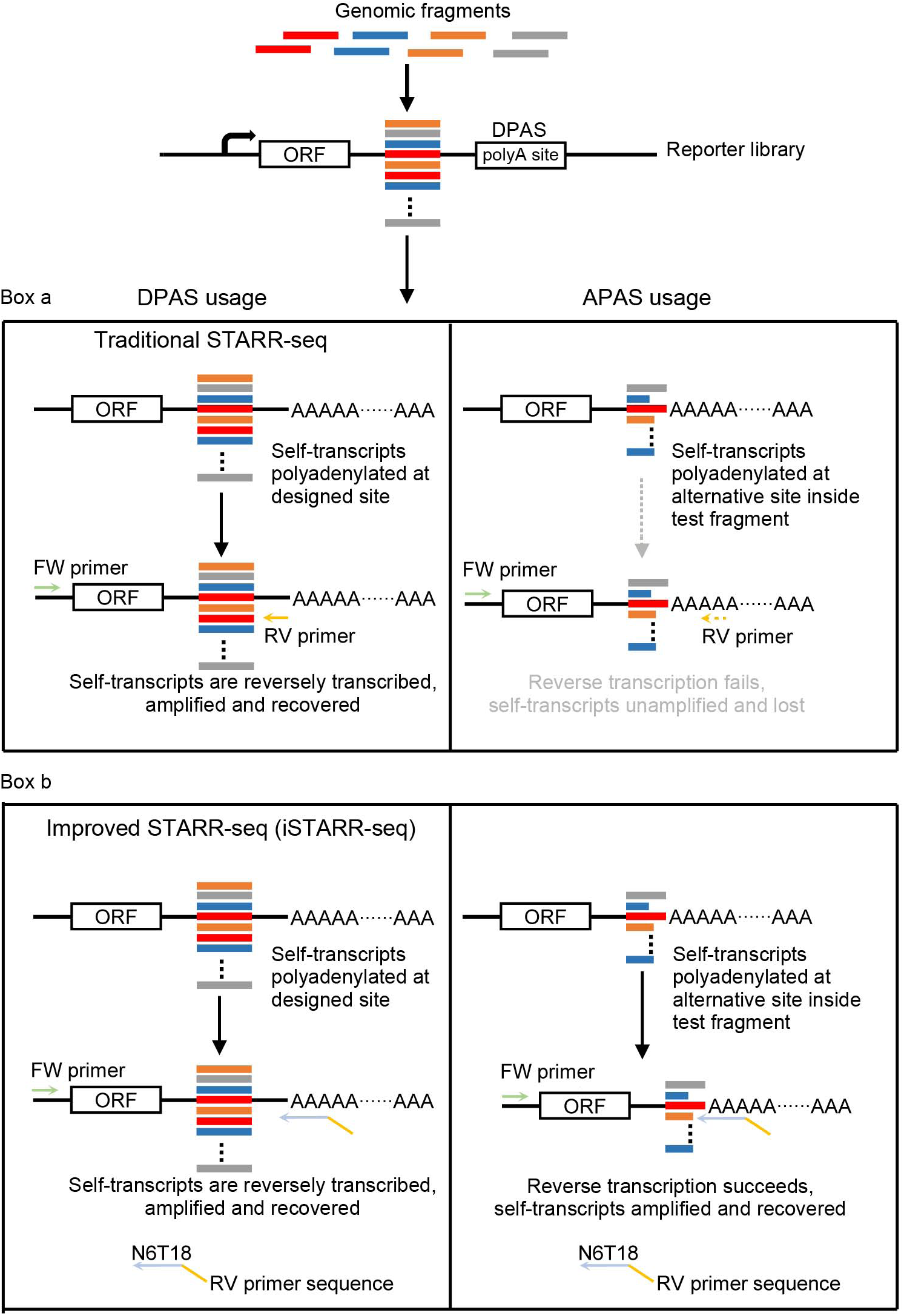
Design of iSTARR-seq to recover lost self-transcripts. DPAS is the pA site shown in the box in the reporter construct, designed to be used exclusively for all self-transcripts from test sequences. APAS is a PAS within the test sequence and can compete with DPAS to be used for the polyadenylation of self-transcripts. Box a shows that in the original design of STARR-seq, if self-transcripts were polyadenylated after the APAS, the sequence between test sequence and DPAS was lost and sequence complementary to the primer used for reverse transcription is located in this lost sequence. This leads to the loss of these self-transcripts in the STARR-seq. Box b shows that a newly designed primer for reverse transcription is used in iSTARR-seq to recover all self-transcripts independent from which PAS used for the polyadenylation.

## Results

### Improved STARR-seq is designed to recover self-transcripts polyadenylated after APAS

Theoretically, only self-transcripts polyadenylated after the DPAS can be detected in the original STARR-seq^18^ (Fig. 1, left side in Box a and b). For this reason, if an enhancer encompass transcription termination site (TTS) and contains a PAS, it is very unlikely to be identified by original STARR-seq because self-transcripts of this sequence are very likely to be polyadenylated after the PAS it contains leading to the loss of downstream sequence complementary to the reverse primer (Fig. 1, right side in Box a). In fact, enhancers identified by STARR-seq are underrepresented at the 3’ end of genes in both *Drosophila* and rice genomes^18, 25^. To avoid the potential loss of these self-transcripts, we redesigned the primer for reverse transcription and named this modified method “improved STARR-seq (iSTARR-seq)” to distinguish it from the original STARR-seq (Fig. 1, Box b). Six random nucleotides and eighteen deoxythymidines were added to the 3’ end of RV, this primer recognizes self-transcripts independent from the PAS usage (Fig. 1, Box b).

### iSTARR-seq recovers self-transcripts polyadenylated after APAS

To find out if self-transcripts were indeed polyadenylated after APAS inside the test DNA sequences and then were lost in STARR-seq, we carried out both STARR-seq and iSTARR-seq experiments in the protoplasts isolated from the leaves of *Arabidopsis thaliana*. Biological replicates for both methods were performed and the sequencing data were highly correlated for plasmid libraries and cDNA libraries derived from self-transcripts (Supplementary Fig. 1a&1b). We combined biological replicates datasets (Supplementary Table 1) for all following analysis. Genome coverage is close to 90% for both methods, and higher in iSTARR-seq (Supplementary Fig. 1c).

The size distributions of the test sequences from STARR-seq and iSTARR-seq plasmid libraries are similar (Fig. 2a) because the plasmids used in transfection were the same. We separated total self-transcripts (tSTs) into two groups based on the difference in the sequence between RV complementary site in the plasmid construct and the test DNA sequence in reporter gene. Self-transcripts polyadenylated after the DPAS are referred as dSTs and self-transcripts polyadenylated after APAS are referred as aSTs. Different from plasmid libraries, considerable amount of self-transcripts recovered by iSTARR-seq are of smaller size (Fig. 2b). Most aSTs are of smaller sizes, while the size range of dSTs is similar to that of the test DNA sequences (Fig. 2c).The aSTs (8.5million) account for about 61% of the tSTs (13.9million) (Supplementary Table 1). We identified 13,424 aSTs enriched regions (Fig. 2d and Supplementary List 1). It is known that APAS are used extensively in the genome of *Arabidopsis thaliana*^26-28^. And most aSTs enriched regions identified by iSTARR-seq overlap with APASs in the genome of *A. thaliana*^26^ (Fig. 2d). These results prove our speculation that the loss of large amount of aSTs occurs because of the use of APAS for the self-transcripts polyadenylation.

**Fig. 2.**
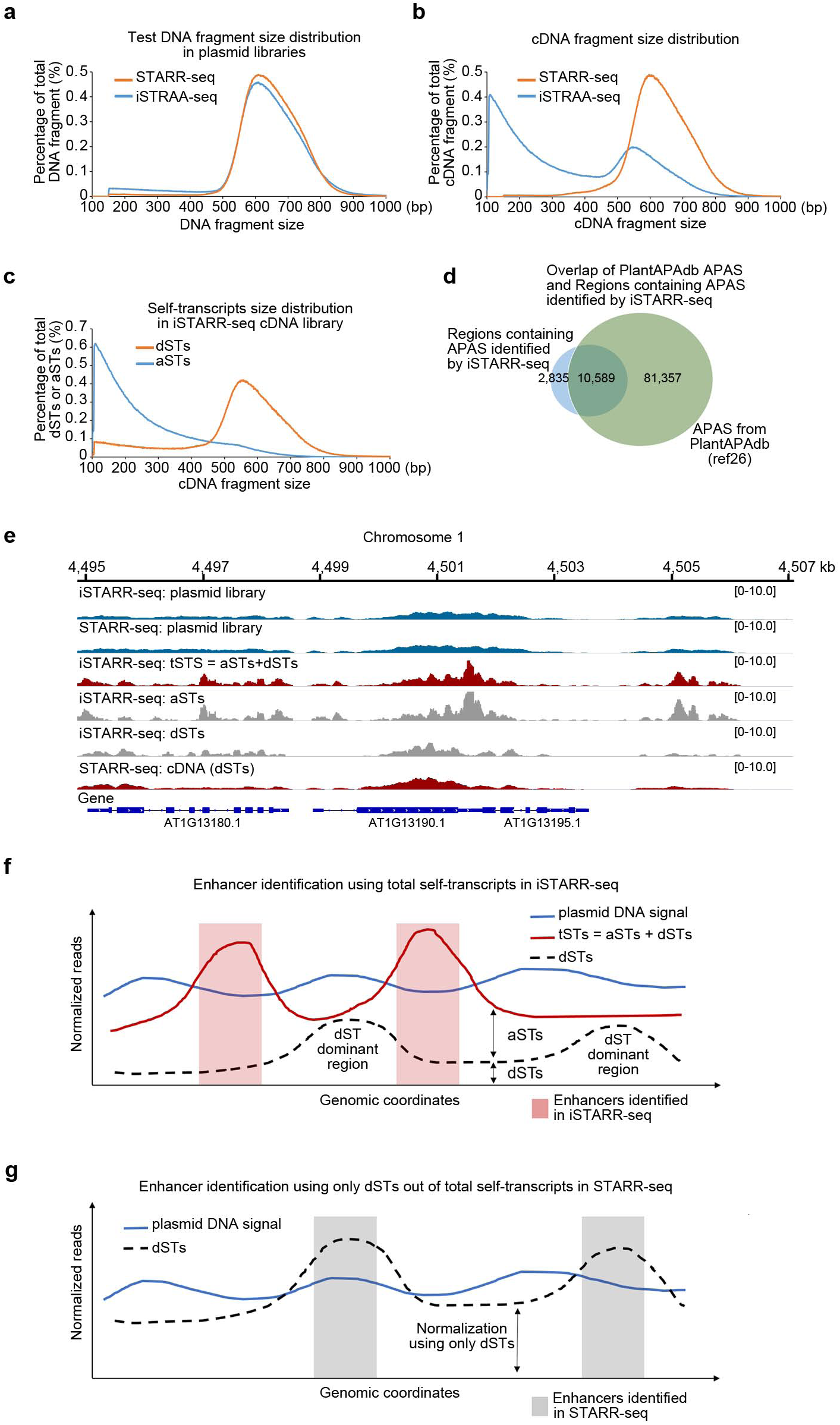
Loss of self-transcripts severely affects enhancer identification. **a**, Size distribution of test fragments from plasmid libraries recovered from transfected protoplasts. **b**, Size distribution of cDNAs in the libraries generated from the self-transcripts recovered from STARR-seq and iSTARR-seq experiments. **c**, Size distribution of self-transcripts polyadenylated after DPAS (dSTs) and APAS (aSTs). **d**, APAS containing regions identified by iSTARR-seq overlap with APAS from PlantAPAdb. **e**, An example region to show STs in iSTARR-seq experiment include both aSTs and dSTs, while the distribution pattern of STs in STARR-seq experiment is similar to the distribution pattern of dSTs from iSTARR-seq experiment. **f**, Simulated normalization of ST and test DNA reads for iSTARR-seq. tSTs include both aSTs and dSTs. Reads of dST dominant regions are not above control after normalization. Regions of low levels of aSTs are named as dST dominant regions. **g**, Simulated normalization of ST and test DNA reads for STARR-seq. Reads in aSTs dominant regions are lost and reads in dSTs dominant regions are biasedly overrepresented.

### Loss of aSTs causes biased signal increase in dST-dominant regions

Original STARR-seq recovers only dSTs but not the aSTs. Visual inspection revealed that the pattern of STARR-seq cDNA signal is more similar to the dSTs from iSTARR-seq (Fig. 2e). The loss of aSTs may cause both local and global effects. To evaluate the potential effects of the loss of aSTs on enhancer identification, we simulated two situations, aSTs are recovered as in iSTARR-seq and aSTs are lost as in STARR-seq (Fig. 2f and 2g). If aSTs were not recovered, the distribution pattern of dSTs shall be different from that of tSTs. For STARR-seq, only reads of dSTs are recovered and are normalized before being compared to the signal of test DNA sequences. In any specific genomic region, the higher is the percentage of aSTs in the tSTs, the less likely the dSTs signal would be increased above the level of the test DNA sequences after global normalization. On the contrary, the lower is the percentage of aSTs in the tSTs, the more likely the dSTs signal will be increased above the level of test DNA sequences in that region after global normalization (2f and 2g). These biases could cause false identification of non-enhancer sequences as enhancers, and at the same time fails to identify real enhancers whose self-transcripts are mostly polyadenylated after APAS.

### Enhancers identified by STARR-seq and iSTARR-seq

By applying the same criteria, we identified 15,862 and 4,956 enhancers for STARR-seq and iSTARR-seq, respectively (Fig. 3a, Supplementary Fig. 2a, b, and Supplementary lists 2 and 3). 2,161 enhancers from this two methods overlap (Fig. 3a), which is significantly higher than their overlap with random sites (*p*<2.2*10^−16^, Fisher exact test). Majority of enhancers identified by STARR-seq (13,701, 86.4%) were not identified by iSTARR-seq. Similarly, 55.8% (2,765/4,956) enhancers identified by iSTARR-seq were absent in STARR-seq (Fig. 3a). The difference in the enhancers identified shall be caused by the loss of aSTs in STARR-seq and the recovery of aSTs in iSTARR-seq. In fact, for iSTARR-seq specific enhancers, the percentage of aSTs in tSTs is significantly higher than enhancers identified by both STARR-seq and iSTARR-seq (Fig. 3b). Also as expected, STARR-seq specific enhancers are of low aSTs signals (Fig. 3b). In two exemplary genomic regions, some enhancers identified by STARR-seq are located in regions of obviously low percentage of aSTs (Enhancers a and b in Supplementary Fig. 3a, and enhancers a, b and d in Supplementary Fig. 3b). And as expected, the loss of aSTs led to the loss of enhancers that were identified by iSTARR-seq (Enhancer ia and ib in Supplementary Fig. 3a and ib in Supplementary Fig. 3b). These results together confirm our speculation that the loss of aSTs changes the signal pattern of self-transcripts across the genome and compromises the reliability of enhancer identification and activity measurement by original STARR-seq.

**Fig. 3.**
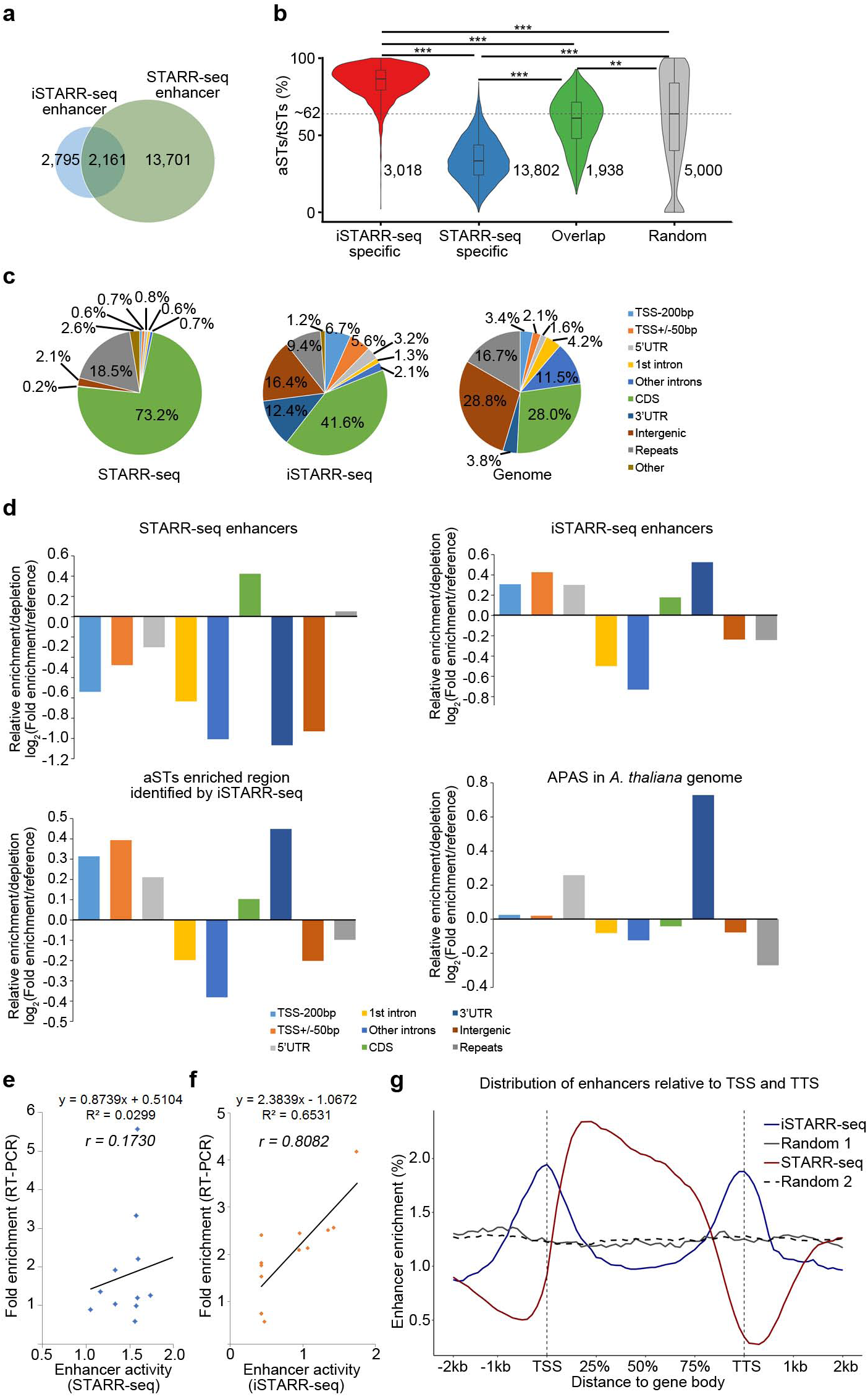
Enhancers identified by STARR-seq and iSTARR-seq. **a**, Overlap between enhancers identified by STARR-sep and iSTARR-seq. **b**, Percentage of aSTs in the tSTs derived corresponding to enhancers identified. ** p<10^−7^, *** p<2.2*10^−16^, Wilcoxon test. **c**, Distribution of enhancers with respect to genomic regions. **d**, Relative enrichment of enhancers and genomic regions containing APAS identified by iSTARR-seq compared to the reference genome. **e**, Randomly selected enhancers identified by STARR-seq were placed in reporter constructs, transfected into same cells, and their enhancing activities were quantified by Real-time PCR. Straight line shows the linear regression. **f**, Randomly selected enhancers identified by iSTARR-seq were placed in reporter constructs, transfected into same cells, and their enhancing activities were quantified by Real-time PCR. Straight line shows the linear regression. **g**, Distribution of enhancers relative to TSS and TTS. Random 1 and random 2 are random genomic sites of the same numbers as iSTARR-seq and STARR-seq enhancers.

The distribution of STARR-seq and iSTARR-seq enhancers in the genome is dramatically different. Most STARR-seq enhancers (73.2%) were mapped to gene coding sequences (CDSs), while much lower percentage of iSTARR-seq enhancers (41.6%) are located in CDSs (Fig. 3c). And the rest of enhancers from STARR-seq and iSTARR-seq are distributed quite differently in all the other genomic regions (Fig. 3c). For STARR-seq enhancers, they are under-represented in all regions except CDS (Fig. 3d), but iSTARR-seq enhancers are over-represented in multiple types of genomic regions, prominently in promoter and terminator regions of genes and less prominent in CDS (Fig. 3d). Consistently, aSTs are similarly enriched in the same genomic regions as iSTARR-seq enhancers and APASs in the *A. thaliana* genome (Fig. 3d). Randomly selected STARR-seq specific enhancers were validated by in vitro transfection and their activities determined by real-time PCR were not well correlated with their activities measured by STARR-seq (Fig. 3e, Pearson’s correlation, r=0.1730). Differently, the activities of iSTARR-seq specific enhancers were nicely correlated with their real-time PCR quantified activities in in vitro test (Fig. 3f, Pearson’s correlation, r=0.8082). These results together show that many STARR-seq identified enhancers could be false positive because of the loss of large amount of aSTs in STARR-seq experiment. At the same time, quite a few enhancers could not be identified as well due to the loss of aSTs. For these reasons, the genetic and epigenetic characteristics of enhancers previously discovered based on STARR-seq identified enhancers can also be misleading. Thus, we decided to use iSTARR-seq identified enhancers for further analysis of the characteristics of enhancers in the genome of *A. thaliana*.

### Enhancers are enriched at both ends of genes

In *Drosophila* and rice genomes, most enhancers mapped by STARR-seq are overrepresented in non-intergenic regions and sequences proximal to TSS, but not at the 3’ end of genes^18, 25^. Similarly, enhancers identified by STARR-seq are strangely overrepresented inside gene bodies in the genome of *A. thaliana* (Fig. 3g). Different from the distribution pattern of STARR-seq enhancers, iSTARR-seq enhancers are overrepresented in two regions around the transcription start sites (TSSs) and the transcription termination sites (TTSs) (Fig. 3g), enriched at both ends of genes in the 5’ and 3’ UTRs (Fig. 3d), and at the same time slightly underrepresented inside gene bodies (Fig. 3g). Many promoters are identified as enhancers in human cells^22^, and it has been suggested that chromatin loop can form between the 5’ and 3’ ends of genes^29^ and regulate transcription in different ways in multiple model organisms^30-37^. And it is also known that the majority of cis-regulatory elements are located close to their target genes in *A. thaliana*^*38*^. Our observation that enhancers are enriched at both ends of genes may also suggest both ends of genes are actively involved in transcription regulation, and possibly through gene looping^30-37^.

### Enhancers are generally accompanying genes of high expression level

Assigning an enhancer to its target gene is challenging. We simplify this task by arbitrarily assuming a gene is a target if the enhancer is located within TSS+/-5kb range of the gene. Total 6,518 genes are assigned as targets for iSTARR-seq enhancers and separated into four groups based on their expression level. The higher is the gene expression level, the higher is the average enhancer number per gene (Fig. 4a). Compared to genes without enhancers in proximity, the expression of these 6,518 genes is significantly higher (Fig. 4b, p < 2.2*10^−16^, Wilcoxon test).

**Fig. 4.**
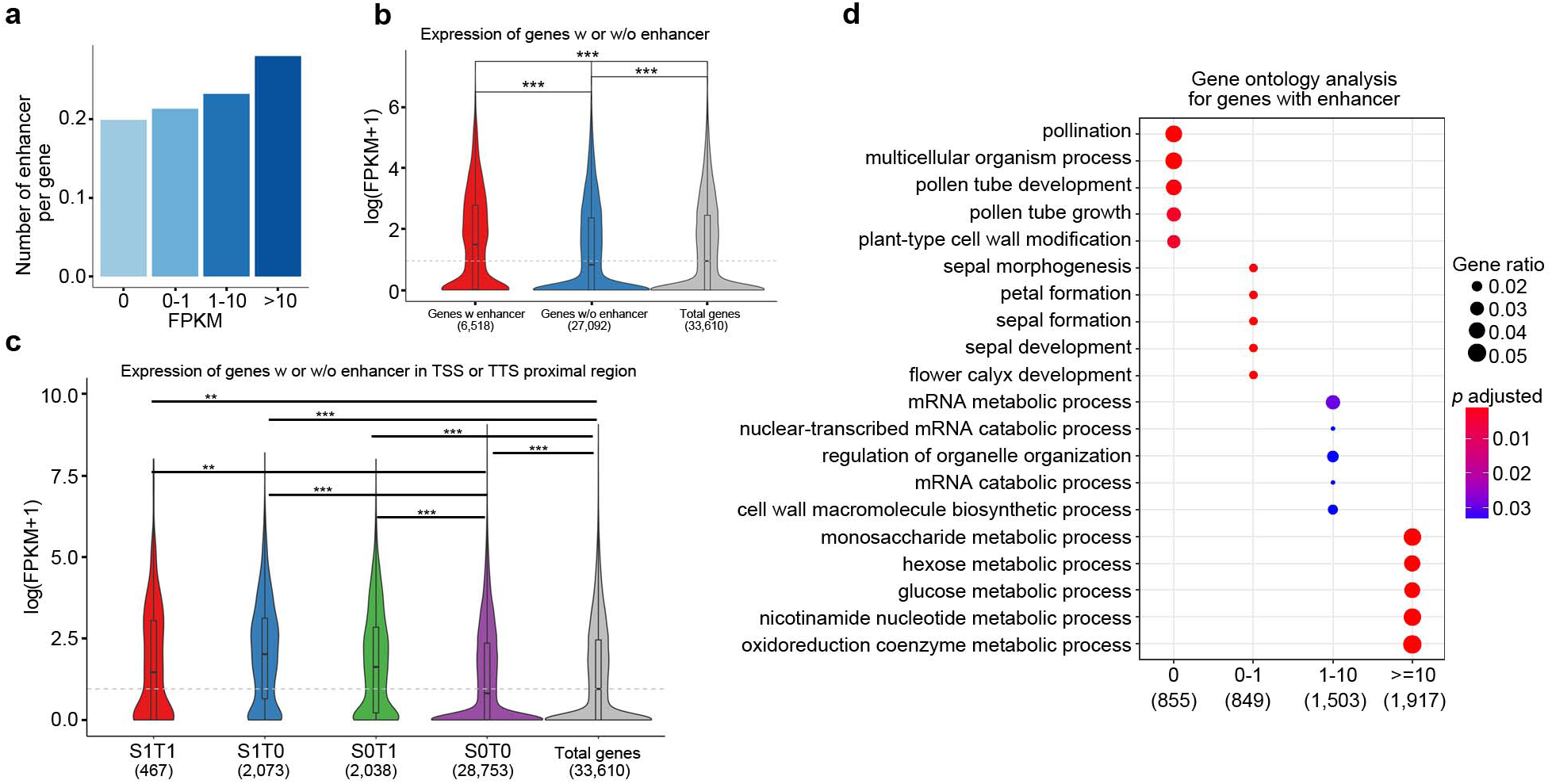
Enhancers activity and gene expression. **a**, For genes of different expression levels, the average number of enhancer per gene was calculated. Enhancers are required to be within 5kb from genes. **b**, Expression levels of genes with or without enhancer in proximity (5kb from TSS) were compared. ***p<10^−10^, Wilcoxon test. **c**, Expression levels of genes with enhancer at TSS or TTS were compared. S1T1 is the group of gene with enhancers at both TSS and TTS ends. S1T0 is the group of gene with enhancer at only TSS end. S0T1 is the group of gene with enhancer at only TTS end. And S0T0 is the group of gene without enhancer at either TSS or TTS ends. **p<10^−7^, ***p<10^−10^, Wilcoxon test. **d**, Gene ontology analysis for genes of different expression level and with enhancer in proximity.

For a specific gene, enhancer can be found at either the TSS or the TTS side (TSS-500bp to TSS+100bp and TTS+/-300bp), or at both sides. We further divided genes with enhancer in proximity into three categories, genes with enhancers at both 5’ TSS and 3’ TTS (S1T1) sides (467), genes with enhancer only at 5’ TSS side (S1T0) (2,073) and genes with enhancer only at 3’ TTS side (S0T1) (2,038). The rest of genes don’t contain enhancer at either ends (S0T0) (28,753). Consistent with previous analysis, genes with enhancer/s are expressed significantly higher levels independent from their location at the 5’ or 3’ end of gene than genes without enhancer (Fig. 4c, p<10^−7^, Wilcoxon test). Expression of genes in groups S1T1 and S1T0 is lower and higher than S0T1 genes, respectively (Fig. 4c), suggesting that the location of enhancers at 5’ or 3’ end of genes may somehow affect the regulation of gene transcription differently.

For genes with proximal enhancers, we further examined if genes of different expression levels are enriched with specific biological functions. Gene ontology (GO) analysis showed that genes expressed at highest level (FPKM>=10) are enriched in metabolic processes (Fig. 4d), genes of medium expression level (1=<FPKM<10) are enriched in mRNA metabolic and catabolic processes (Fig. 4d). These genes are generally house-keeping genes and their functions are ubiquitously required for different types of cells. Different from genes expressed at medium and high levels, genes expressed at lowest level or silent (0=<FPKM<1) are enriched in development and reproduction processes, such as pollination, pollen tube development and growth (Fig. 4d). These preliminary results suggest that the functions and characteristics of enhancer could be quite diverse depending on their location in the genome and the genes that they regulate.

### Epigenetic clustering reveals four types of enhancers in *A. thaliana*

Enhancers identified by iSTARR-seq may carry different epigenetic marks at their endogenous genomic loci. To characterize the epigenetic states of enhancers, we collected datasets of chromatin accessibility, RNAPII binding, DNA methylation and histone modifications in the leaf cells of *A. thaliana*^39-48^(Supplementary list 4). Enhancers are ranked by the overall signal intensity of multiple epigenetic features (Fig. 5a). Consistent with the observation in *Drosophila*^*18*^, enhancers are mostly accessible to the digestion by enzymes of DNaseI and Tn5, and enriched with RNAPII (Fig. 5a). Also consistent with our previous observation in rice^25^, H3K4me1 is largely absent from enhancers in *A. thaliana*, while H3K4me3 is enriched at enhancers (Fig. 5a). H3K27ac is another mark frequently used to predict active enhancers in mammalian cells^17^, and it is also enriched but together with other histone acetylation marks like H3K9ac, H3K36ac and H3K56ac (Fig. 5a), supporting a previous claim that H3K27ac may not be a unique mark for plant enhancers in the flower of *A. thaliana*^49^.

**Fig. 5.**
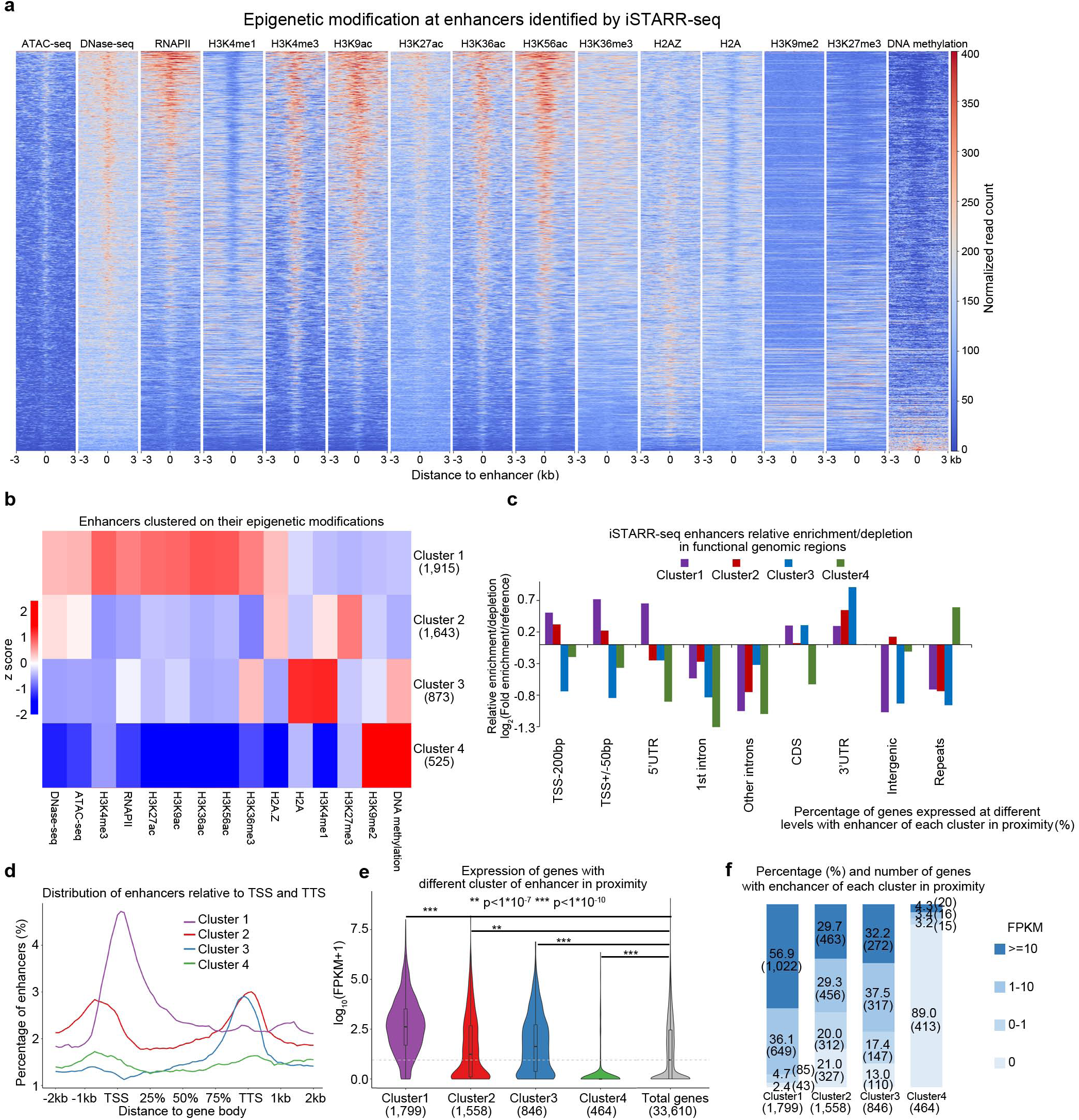
Regulation of enhancer activity at endogenous chromatin. **a**, iSTARR-seq identified enhancers are arranged from top to bottom according to the overall intensity of epigenetic signals at enhancers. **b**, iSTARR-seq identified enhancers are clustered into four groups on their epigenetic characteristics. **c**, Relative enrichment of depletion of the four clusters of iSTARR-seq enhancers in functional genomic regions. **d**, Distribution of the four clusters of iSTARR-seq enhancers relative to the TSS and TTS. **e**, Expression levels of genes with enhancers of different cluster in proximity. **p<10^−7^, ***p<10^−10^, Wilcoxon test. **f**, The expression levels of genes associating with enhancers of different cluster. Gene number is shown in parentheses.

We further clustered all enhancers into four categories (Fig. 5b and Supplementary Fig. 4). Cluster 1 is the largest group containing 1,915 enhancers that are accessible to DNaseI/Tn5 digestion, enriched with nearly all positive histone modifications (except H3K4me1), and are absent of repressive marks (DNA methylation, H3K9me2 and H3K27me3) (Fig. 5b). Enhancers in cluster 2 (1,643) are also accessible to DNaseI/Tn5 but at lower rates and enriched with H3K27me3, as well as low levels of H3K4me1 and H2A.Z (Fig. 5b). Cluster 2 enhancers may be in poised state at their endogenous genomic loci. Enhancers in cluster 3 (873) are less accessible to digestion (Fig. 5b). However, these enhancers carry medium level of H3K36me3 and high level of H3K4me1, and also medium DNA methylation which may correlate with either active or repressive chromatin types dependent on their locations in the genome and their chromatin context. Enhancers in cluster 4 (525) are not accessible to DNaseI/Tn5 digestion and are enriched with repressive H3K9me2 and DNA methylation (Fig. 5b). These enhancers may only function during cell type-specific developmental or differentiation stages. The four categories of enhancers also show strikingly different distribution patterns in the analyzed genomic regions (Fig. 5c) and different distances relative to TSS and TTS (Fig. 5d).

The different epigenetic states of the four categories of enhancers may correlate with their functions at their endogenous genomic loci. To test this hypothesis, we assigned the nearest genes to the enhancers of each cluster and compared the levels of gene expression. As expected, genes (1,799) associating with cluster 1 enhancers are expressed at highest levels and genes (464) associating with cluster 4 enhancers are mostly silent (Fig. 5e). Expression of genes (1,558) associating with cluster 2 enhancers is slightly higher than control (Fig. 5e), suggesting these group of enhancers may be in a poised state associating with low level of gene expression, and H3K27me3 enriched chromatin is facultative heterochromatin subjective to regulation. Genes (873) associating with cluster 3 enhancers are expressed at levels higher than genes associating with cluster 2 enhancers (Fig. 5e). Cluster 3 enhancers are enriched at the 3’ end of genes and carry unique epigenetic modifications (DNA methylation, H3K4me1 and H3K36me3). These results suggest that cluster 3 enhancers may function through mechanism that is different from cluster 1 enhancers.

Genes of different expression levels were further analyzed and were associated with different types of enhancers. Genes of low or no expression (0=<FPKM<1) are enriched in development processes (Fig. 4f). There are 639, 257 and 428 genes (41%, 30.4% and 92.2%) associating with enhancers of clusters 2, 3 and 4 that are expressed within this range (Fig. 5f), while only 128 genes (7.1%) associating with cluster 1 enhancers are expressed in this range (Fig. 5f), suggesting enhancers of clusters 2, 3 and 4 are more involved in developmental gene regulation than enhancers of cluster 1. Genes of medium expression (1=<FPKM<10) are enriched in catabolic processes (Fig. 4f). Total 649, 456 and 317 genes (36.1%, 29.3% and 37.5%) are associating with cluster 1, 2 and 3 enhancers, only 16 genes (3.4%) associating with cluster 4 enhancers are expressed in this range (Fig. 5f). Genes of highest expression level (FPKM>=10) are mostly house-keeping genes (Fig. 4f). Total 1,022, 463 and 272 genes (56.9%, 29.7% and 32.2%) associating with enhancers of clusters 1, 2 and 3, while only 20 genes (4.3%) associating with cluster 4 enhancers are expressed at this level (Fig. 5f). These results show that enhancers of cluster 1 are mostly regulating highly expressed genes. Enhancers of clusters 2 and 3 seem to be more functionally diversified and regulating genes ranging from lowest to highest levels. And enhancers of cluster 4 are predominantly co-localizing with genes of lowest expression levels.

### Transcription factor binding sites are enriched differently in the four clusters of enhancers

Enhancer DNA provides a platform for transcription factors to bind^1^. Transcription factor binding sites (TFBS) across the genome of *A. thaliana* had been determined by DAP-seq^50^. For iSTARR-seq enhancers, 28 families of TFBSs are enriched (Fig. 6a, blue dots). Though many families of TFBSs are also enriched in STARR-seq enhancers (Fig. 6a, orange dots), nearly all of them are enriched at a lower level (Fig. 6a). For the four clusters of enhancers, the enrichment of TFBS varies dramatically (Fig. 6b-e). TFBSs are most conservatively enriched at clusters 1 and 2 enhancers (Fig. 6b,c), even though cluster 2 enhancers may be mostly in poised state and are not associating with very high levels of gene expression (Fig. 5e,f). Surprisingly, majority of TFBS are absent from clusters 3 and 4 (Fig. 6d, e). Cluster 3 enhancers are well correlated with decent high level of gene expression (Fig. 5d) but lack of most active histone modifications (Fig. 5b), these suggest that cluster 3 enhancers may function through a not well characterized mechanism, or possibly through gene looping, a three-dimensional structure that efficiently facilitates the recycled use of RNAPII^37^. Similar to cluster 3 enhancers, cluster 4 enhancers are not enriched with most TFBS examined (Fig. 6e). Cluster 4 enhancers are overrepresented in repetitive sequences (Fig. 5c) in which known transcription factor binding sites may not as enriched as in non-repetitive sequences. Another possibility is that many development- or differentiation-specific TFBSs have not been well-characterized thus were not analyzed in this study. In sum, iSTARR-seq identified enhancers are very diverse in their potential for the binding of different categories of transcription factors. These differences are encoded in the DNA sequences that determine which transcription factors to recruit. I addition, the temporal- and spatial-specific expression of transcription factors adds another layer of complexity to the complex mechanisms controlling gene expression during differentiation and development.

**Fig. 6.**
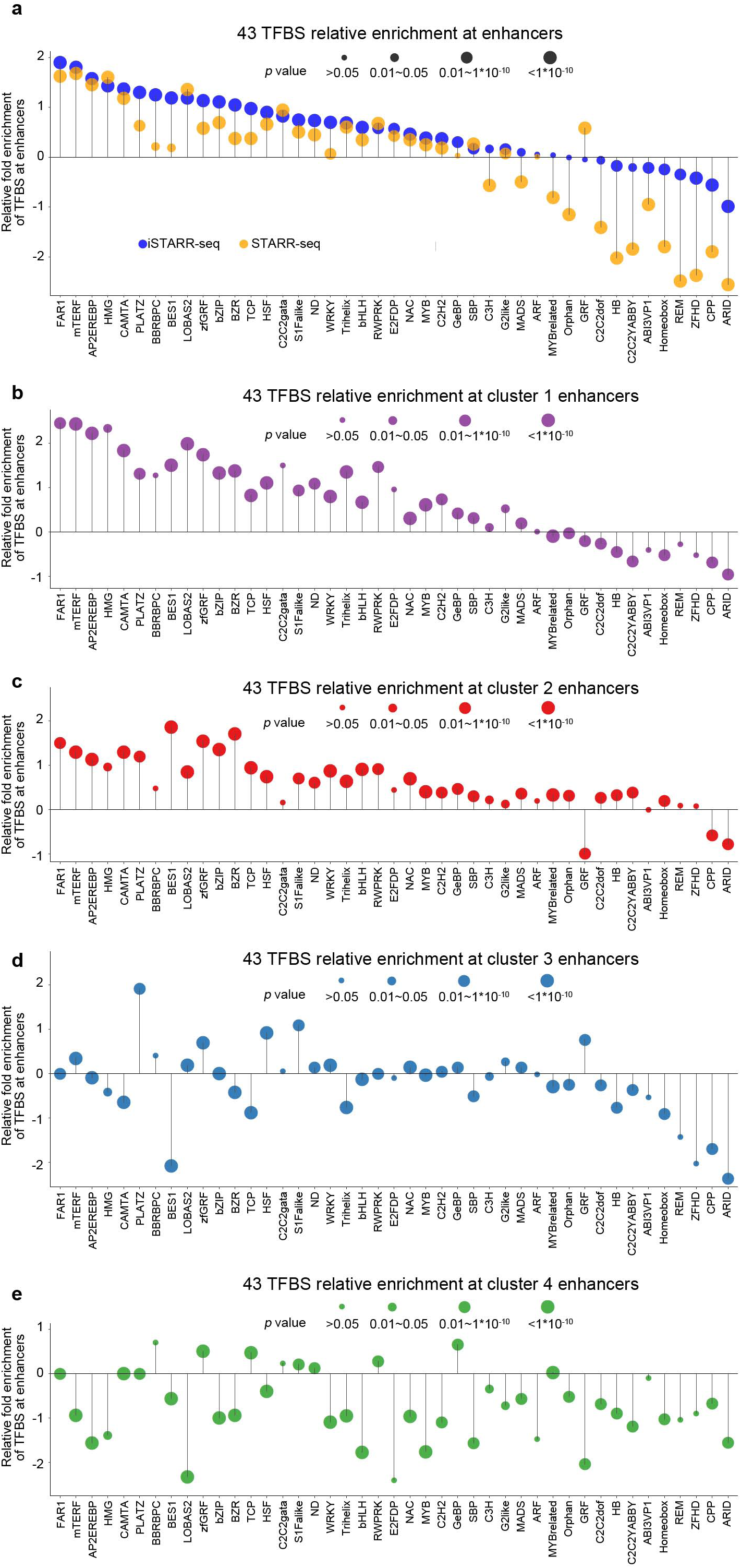
Enrichment of transcription factor binding sites at enhancers. **a-e**,TFBS enrichment or depletion relative to random control sites was determined for iSTARR-seq and STARR-seq enhancers **(a)**, for four clusters of iSTARR-seq enhancers **(b-e)**. P values were calculated by comparing the enrichment of TFBSs in enhancers and random control sites with Fisher exact test.

## Discussion

Cis-regulatory elements in the genome of *A. thaliana* are predominantly located close to genes^29, 38^. Contrary to this, methods frequently used to predict enhancers arbitrarily exclude DHSs close or within genes and keep only DHSs distant from genes as enhancer candidates^46, 49^. Histone marks that are being used to predict enhancers in animals have been questioned if they are ideal enhancer marks in plants^25, 49^. STARR-seq is one of many methods that had been developed to identify functional enhancers^18^. Here we show that APAS in the test DNA sequences can be used for polyadenylation, and self-transcripts polyadenylated after APAS cannot be recovered using the reverse primer of STARR-seq. APASs usage is common in *A. thaliana*^*27, 28*^. The effect of the loss of aSTs on enhancer identification is obvious as shown in this work. This has important implications for methods derived from STARR-seq and methods based on quantification of self-transcripts. Those methods have become increasingly more important for global regulatory elements identification efforts of large consortia and especially for the new stage of ENCODE projects emphasizing on functional analysis^51^, and new prediction method^52^ that had been developed based on enhancers identified by STARR-seq could also be affected.

The percentage of tested fragments containing APAS in a specific genome may vary dramatically. Thus, that how severe is the effect of the loss of aSTs on the enhancer identification and activity quantification depends on the unique sequence composition of each different genome that carries different number and distribution pattern of APAS. Based on our preliminary analysis, we find aSTs account for 20%-30% of total self-transcripts for *Drosophila* and rice genomes, and ∼10% for human genome (ongoing projects, not published). These results confirm again that the recovery of aSTs is important for reliable characterization and activity quantification of enhancers.

Besides, we discovered an additional application for iSTARR-seq that it can be used to identify APASs. We identified 13,424 regions containing APAS in the genome of *A. thaliana* (Supplementary List 1) and most of them can be found in the PlantAPAdb^26^ (Fig. 2d), confirming indeed iSTARR-seq can be used for APAS identification.

The loss of aST signal frequently occurs at genomic sequences containing PAS. As revealed by this work, enhancers are actually also enriched at the 3’ end of genes (Fig. 3c,d,g), dramatically different from enhancer distribution patterns in rice and *Drosophila*^*18, 25*^ and also STARR-seq enhancers identified in this work. In fact, looping between the promoter and terminator regions of a gene had long been proposed as a mechanism regulating gene transcription ^29, 32, 33, 35-37^. We used published chromatin loop data^29^ and carried out chromatin interaction analysis using iSTARR-seq enhancers as anchors and found no significant higher level of long-range enhancer-promoter interactions (data not shown). Most genes in the genome of *A. thaliana* are short and the chromatin interaction data we used were generated by traditional Hi-C which biasedly reduces chromatin interaction frequency within short genomic distance^53^. Higher resolution and bias-free Hi-C method like SAFE Hi-C^53^ can be used and may possibly reveal the finer chromatin loops anchored at enhancers.

Epigenetic marks that are being used to predict enhancers in plants are lack. Based on the four clusters of enhancers identified by iSTARR-seq, we propose to use combinations of multiple epigenetic marks, instead one or two histone modifications, to predict if a DNA element in the endogenous genomic locus is a specific type of enhancer. For example, if a DNA sequence is accessible, bound by RNAPII, carrying H3K4me3, H3K36me3, and H3K9ac, H3K27ac, H3K56ac and flanked by H3K36ac, but not enriched with H3K4me1, H3K9me2, H3K27me3 and DNA methylation, this site could very likely be an active enhancer of the cluster 1. Similarly, multiple epigenetic marks can also be used in combination to predict other types of enhancers. How successful this approach can be is subjected to more tests. And similar work has been recently published based on enhancers identified in *Drosophila* by STARR-seq^*52*^.

In sum, we present here a solution to solve a systematic flaw in STARR-seq, a method used widely now days for functional enhancer identification, and even being modified for silencer identification in mammalian genomes^54^. Resolving this systematic error is important for increasingly intensive efforts to identify all kinds of gene regulatory elements in the genomes of model organisms, and to understand how gene transcription is precisely controlled in both ubiquitous and cell type-specific ways during development and differentiation.

## Materials and methods

### Screening vector

For STARR-seq in the cells of *A. thaliana*, we constructed a screening vector based on the plasmid pPBI221 backbone^55^ and modified this vector by introducing several sequences, which include a CMV 35S mini promoter ^46^, an intron and a GFP sequence, which are arranged sequentially and are underlined in the following DNA sequence.

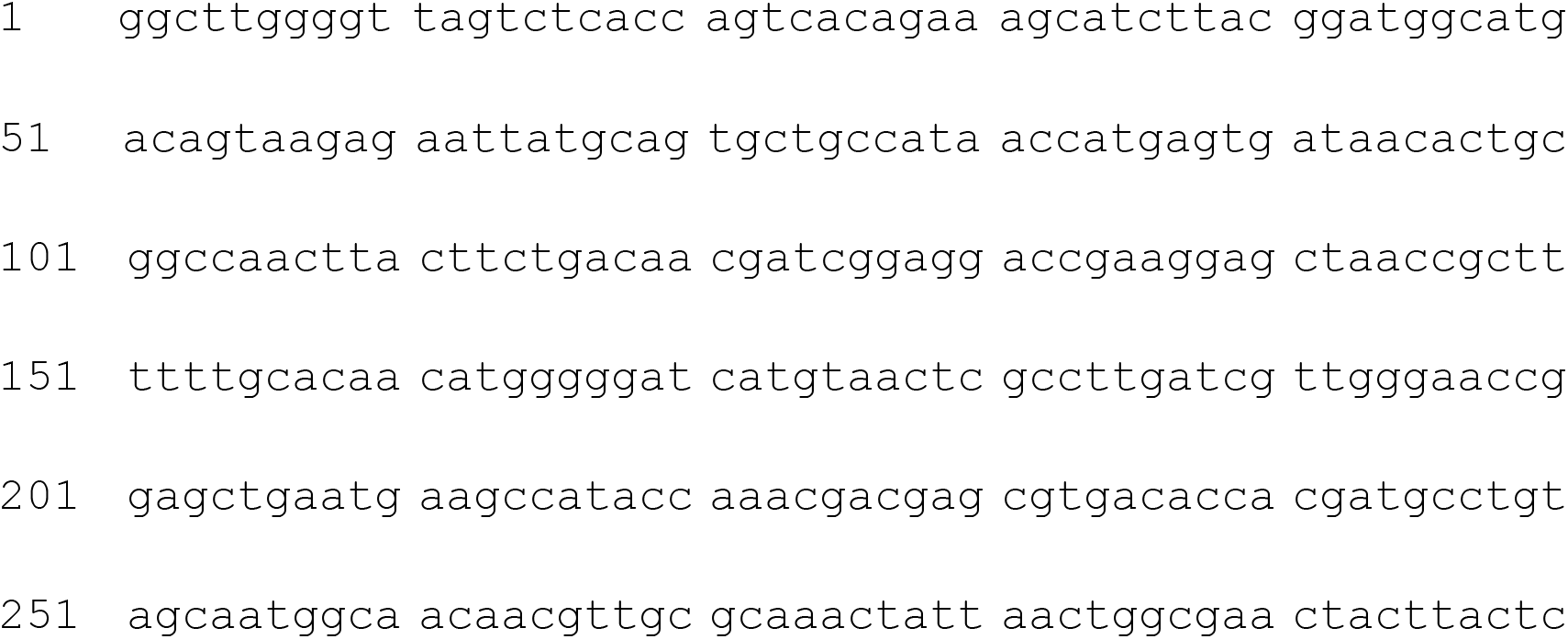
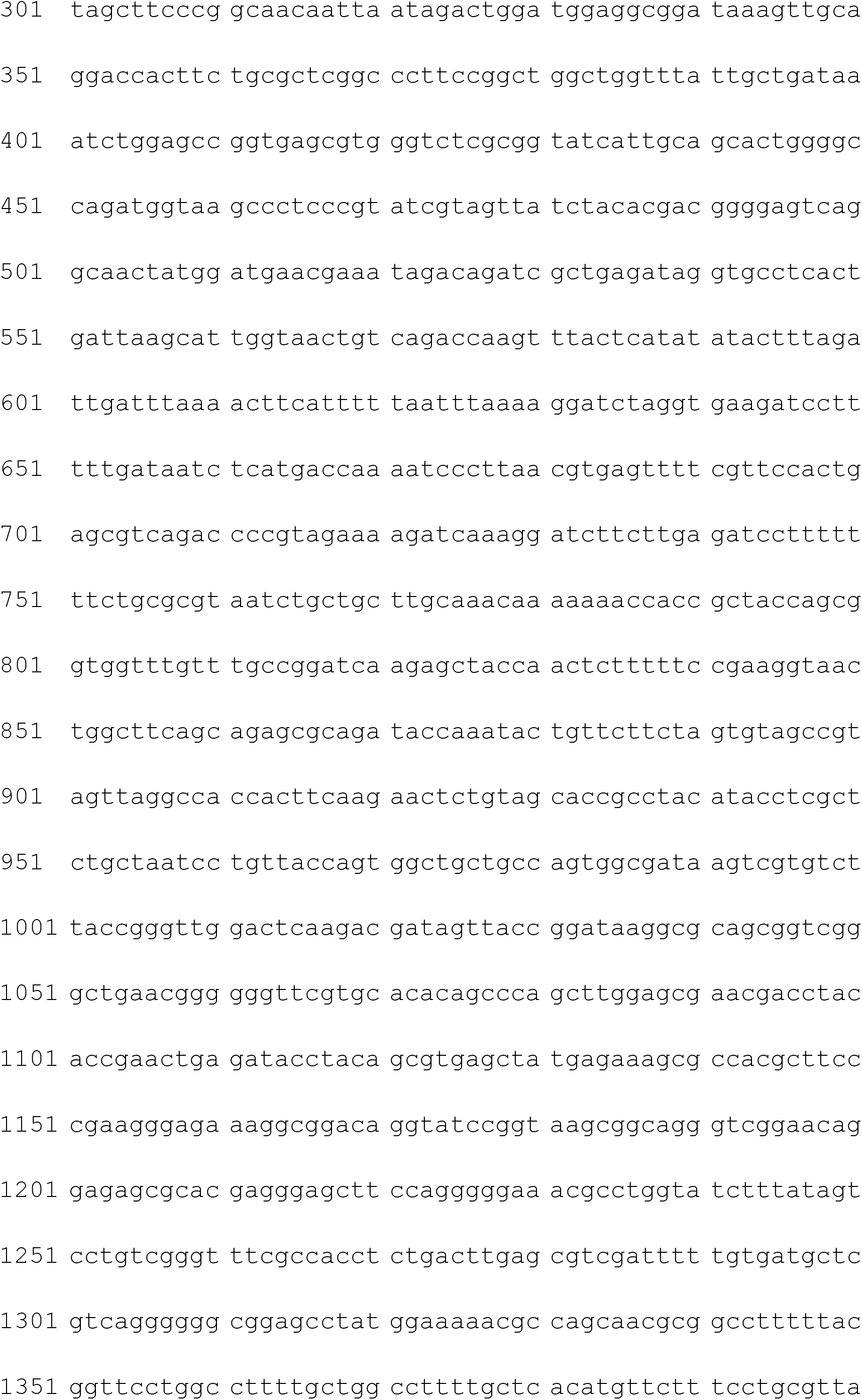
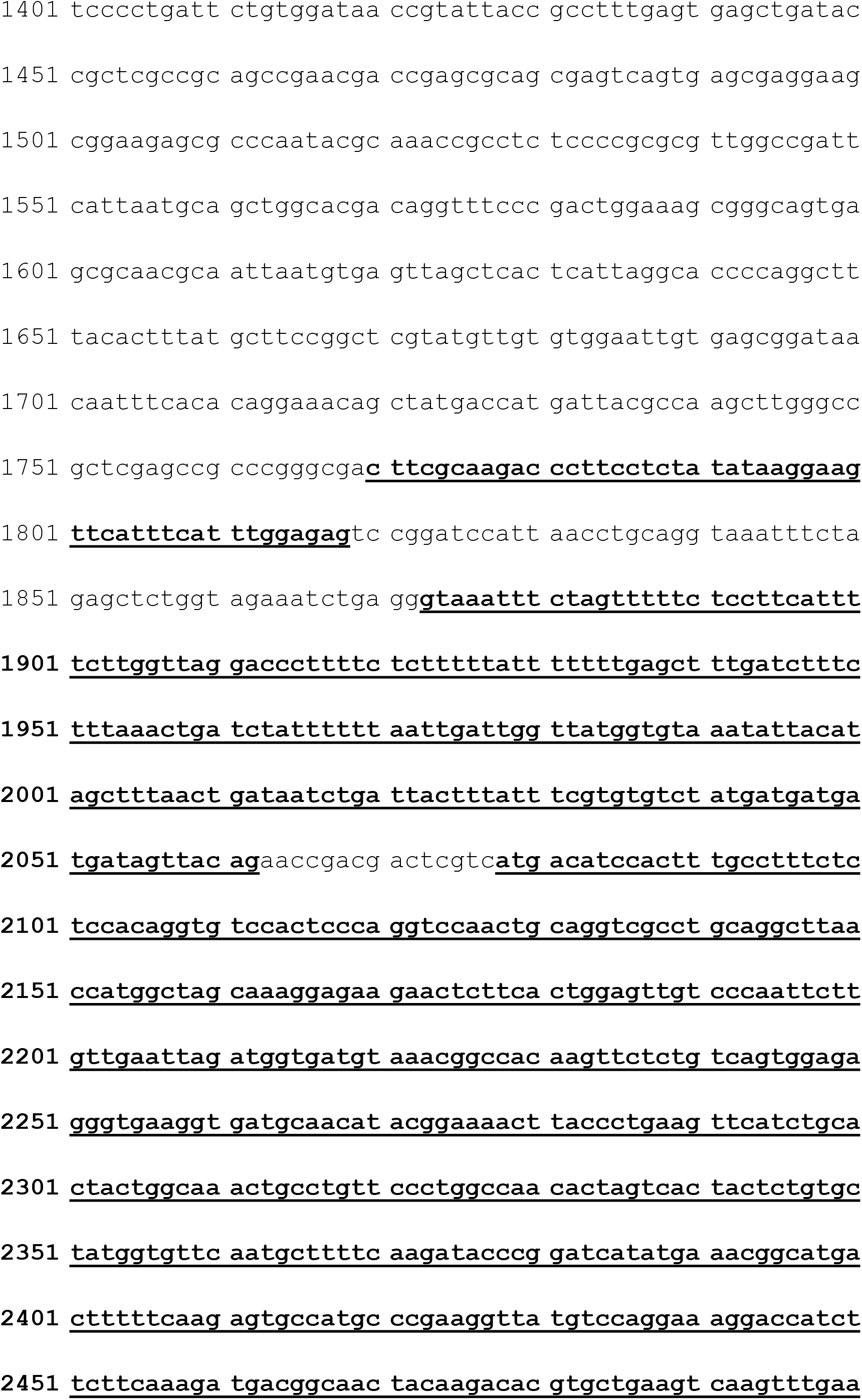
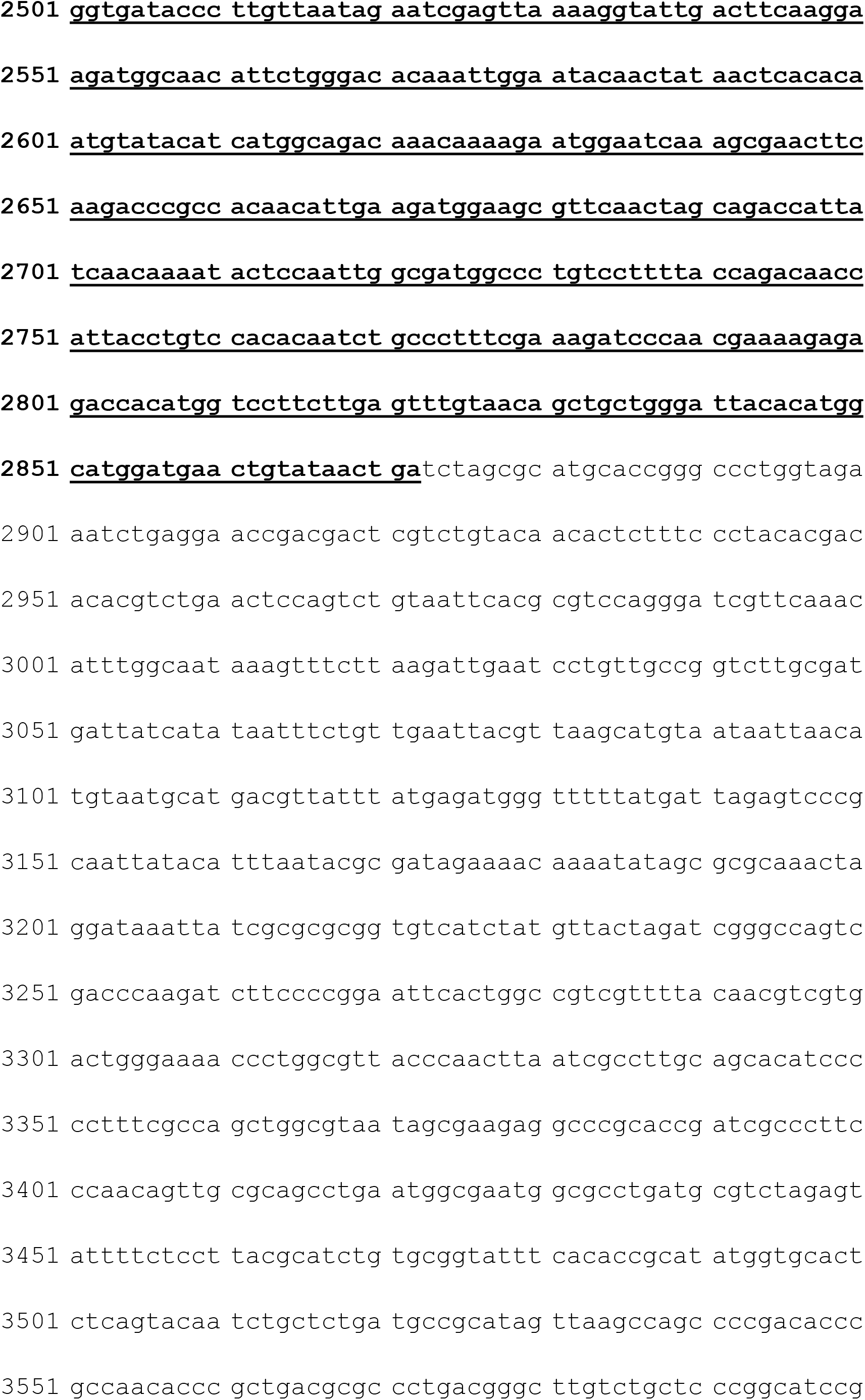
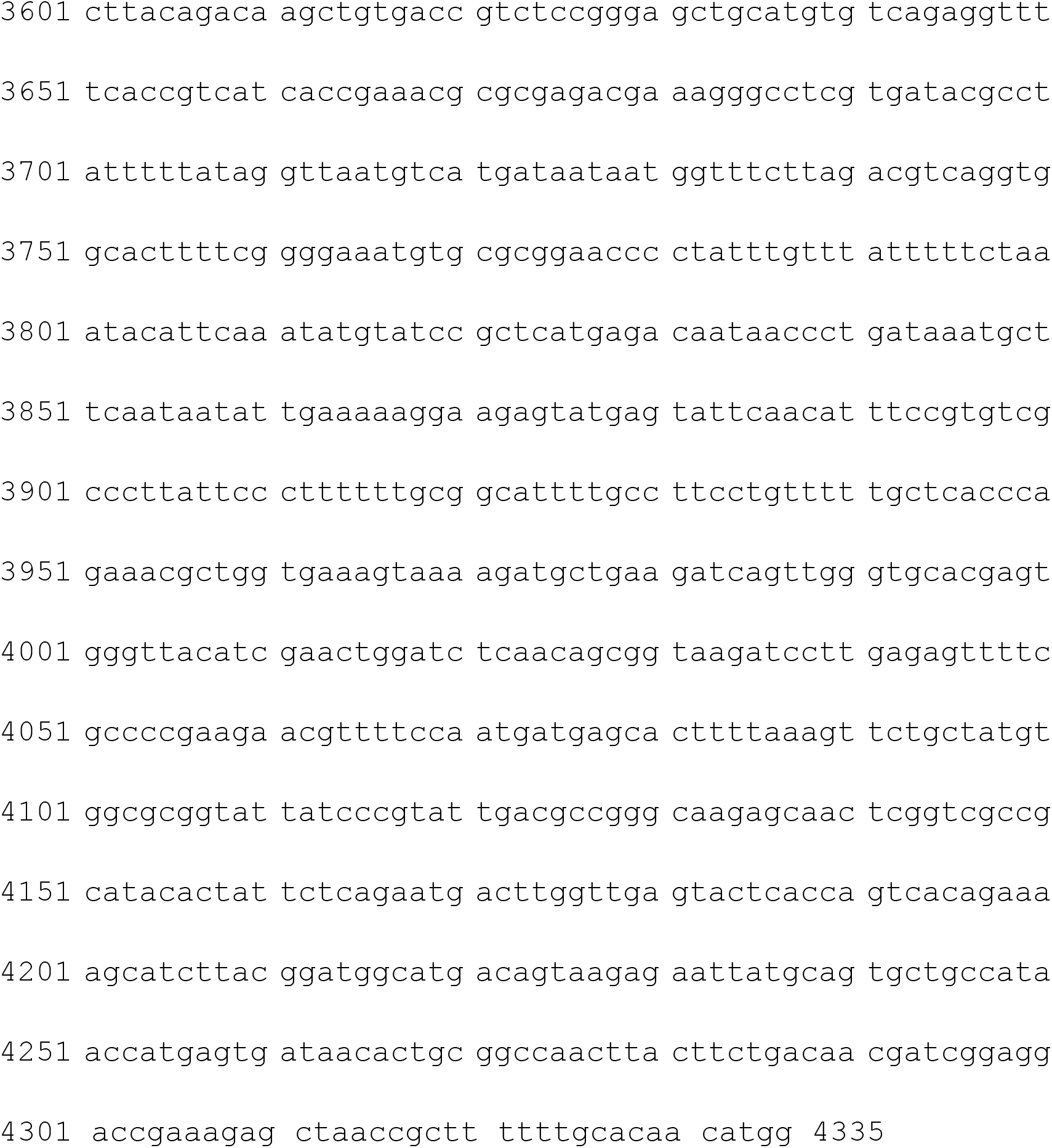

### Generation of input (screening) libraries

We extracted genomic DNA from the seedlings of *A. thaliana* of two weeks. About 30µg of genomic DNA was diluted to 50ng/µl and fragmented by sonication (Scientz II-D). DNA fragments (500bp-800bp) were selected, e nd repaired, 5’-phosphorylated and 3’ dA-tailed with VAHTS Mate Pair Library Prep Kit for Illumina® (VAHTS; cat. no. ND104). VAHTS Adapters for Illumina was ligated to about 6µg of DNA fragments using VAHTS Mate Pair Library Prep Kit for Illumina® (VAHTS; cat. no. ND104) following the manufacturer’s protocol. Adaptor ligated DNA was purified by using GeneJET PCR Purification Kit (Thermo Scientific; cat. no. K0702) then was amplified by TransStart FastPfu Fly DNA Polymerase (Transgen; cat. no. AP231) with Illumina sequencing primers

Forward primer: ACACT CTTTC CCTAC ACGAC G

Reverse primer: GACTG GAGTT CAGAC GTGTG C.

We obtained linear pPBI221 by PCR amplification (95°C for 5min; then 25 cycles of 95°C for 20s, 57°C for 20s and 72°C for 4min;

Forward primer: ACACG TCTGA ACTCC AGTCT GTAAT TC

Reverse primer: GTCGT GTAGG GAAAG AGTGT TGTAC A.

Circular plasmids was removed by digestion with DpnI (NEB; cat. no. R0176) at 37°C for 30min. PCR products were recovered with E.Z.N.A.® Gel Extraction Kit (Omega; cat. no. D2500). The adaptor ligated genomic DNA were recombined to linearized screening vector with ClonExprress II One Step Cloning Kit (Vazyme; cat. no. C112). For each reaction, 100ng linearized vector and 40ng adaptor ligated DNA was used in a total volume of 20µl. 70 reactions were performed in total. Each of 140 tubes (100µl each) of Trans1-T1 Phage Resistant Chemically Competent Cell (Transgen; cat. no. CD501) was transformed with 10µl of DNA, according to the manufacturer’s protocol. 140 transformation reactions were pooled, transferred to 4L LB_AMP_ medium, and grown to OD 0.8. Plasmids were purified using E.Z.N.A.® Endo-Free Plasmid Maxi Kit (Omega; cat. no. D6926) then quantified.

### Protoplast preparation

The *Arabidopsis* seedlings were grown at 22°C with a relative humidity around 60%. The light/dark photoperiod is 16/8 h by using a bank of 400 W metal halide lamps. The light intensity is 400µmol m^−2^s^−1^ at the plant height. We collected 2-week-old seedlings for the all experiments. Protoplasts isolation was performed following previously published protocols^56^. Leaves tissues from 2-week-old seedlings were used. Seedlings were cut together into approximately 0.5 mm strips using sharp razors in 0.6M D-mannitol on Petri dishes. The strips were then digested (1.5% Cellulase RS, 0.6M mannitol, 0.1% BSA, 0.75% Macerozyme R-10, 3.4mM CaCl2, 10mM MES, pH 5.7), and then vacuumed at −50kPa in bottle for 30 min. After that, the bottle was put on oscillator for 4-5h in dark with gentle shaking (45rpm) at 22°C. After digestion, protoplasts were filtered through 300 micron nylon meshes and washed 3-5 times with W5 solution (125mM CaCl2, 154mM NaCl, 5mM KCl, 2mM MES, pH5.7). Protoplasts were pelleted by centrifugation at 80g for 5min. After another two wash with W5 solution (equal volume), protoplasts were then resuspended in MMG solution (15mM MgCl2, 0.6M mannitol, 4mM MES, pH5.7), and placed on ice for 30 min. Protoplasts were centrifuged at 80g again for 5 min and finally resuspended at a concentration of 1×10^7^ cells per ml using the MMG solution, determined by using a hematocytometer. All procedures were carried out at room temperature.

### Plasmid library transformation

Protoplasts transient transformation was carried out as described^56^. For each sample, 30-40μg of plasmid was mixed with 100μL protoplasts (∼1×10^6^ cells) in a 2ml tube with 110μl of freshly prepared PEG solution [40% (W/V), 0.1M CaCl_2_, 0.6M mannitol] added. Gently mix, then incubate in dark at 28°C for 15 min. Add 800μl of W5 solution slowly. Invert gently to mix well, then centrifuge immediately at 80g for 5 min. The pelleted protoplasts were resuspended gently in 800μl of W5 solution. Finally, protoplasts transfected were cultured at 22°C for 24h in dark. Reporter GFP signals were checked under fluorescent microscopy.

### Reporter cDNA libraries construction for Illumina sequencing

24hrs post-transfection, cells were washed in W5 solution twice and concentrated. Total RNA was extracted using TransZolTM Up Plus RNA Kit (ER501). polyA+ RNA fraction was isolated using VAHTS mRNA Capture Beads (N401). 5U DNase I (NEB M0303S) was used to digest DNA at 37°C for 20 min.

Synthesis of first strand cDNA was carried out with TransScript One-Step gDNA Removal and cDNA synthesis SuperMix (AU311 50°C for 30min and 85°C for 5s) using a specific primer N6T18RV (5’ GACTG GAGTT CAGAC GTGTG CTCTT CCGAT CTttt ttttt ttttt ttttt NNNNN N 3’) and 8ul of RNA in 20 reactions. All reactions were pooled. The total amount of reporter cDNA was amplified for Illumina sequencing by a 2-step nested PCR strategy using the TransStart® FastPfu Fly DNA Polymerase (AP231-12).

### First round PCR amplification and Cas9 treatment

In the first round PCR (98°C, 2min; then 10- 15 cycles of 98 °C for 30s, 58°C for 30s, 72°C for 30s; and then 72°C for 5min), 10-30ng cDNA were used for amplification with 2 reporter-specific primers (Forward primer: 5’Biotin GACTG GAGTT CAGAC GTGTG C3’ & Reverse primer T18RV: 5’ GACTG GAGTT CAGAC GTGTG CTCTT CCGAT CTttt ttttt ttttt ttttt 3’), one of which spans the RNA splice junction of GFP intron. PCR products were purified by GeneJET PCR Purification Kit (K0701) and eluted in 30ul 1XCas9 Buffer. And then the PCR products were cut by Cas9. sgRNA transcription by ThemorFisher Kit (TranscriptAid T7 High Yield Transcription Kit, Thermo Scientific). Nuclease-free H_2_O 23ul, Cas9 Nuclease Reaction Buffer (10×) 3ul, sgRNA 3ul (300nM), Cas9 Nuclease 1ul. Incubate at 37°C for 10min before the adding of the PCR products and incubation at 37°C for another 60min. The PCR products were then captured by magnetic streptavidin C1 beads. After Cas9 treatment, the extended products were captured by pre-washed magnetic streptavidin C1 beads (Invitrogen, 650.01) in 1×Binding & Wash (B&W) buffer (10mM Tris-HCl pH 8.0, 0.5mM EDTA, 1M NaCl) in a 1.5ml Eppendorf Lobind tube (Eppendorf, 0030 108.051). DNA binding was performed in a thermo mixer (Eppendorf, 5355 000.011) for 30 min by mixing at 1400 rpm (10s on, 10s off) at 23°C.Then, the beads were washed once with100ul of 1×B&W buffer, three times with 150ul of EBT buffer (10mM Tris-HCl pH 8.0, 0.02% Triton X-100) and resuspended in 8.4ul of elution buffer (EB; 10mM Tris-HCl pH8.0) in preparation for PCR.

### Second round PCR amplification to generate cDNA sequencing library

Purified PCR product as template for the second round PCR (98°C for 2min; followed by 6-10 cycles of 98°C for 30s, 65°C for 30s, 72°C for 30s; and then 72°C for 5min) with TransGen® Biotech FastPfu Fly DNA Polymerase (AP231) and with VAHTS™ DNA Adapters for Illumina® (N302). PCR products were purified by GeneJET PCR Purification and eluted in 20-30ul EB.

### Transfected plasmid library construction for Illumina sequencing

When the PolyA RNA was captured, keep the total RNA (without mRNA) supernatant, add 10ul RNase A TransGen® Biotech GE 101 20mg/ml at 37°C for 30-60min, and the plasmid was purified by GeneJET PCR Purification Kit and eluted in 50ul EB. Purified plasmid as template for the second round PCR (98°C for 2min; followed by 8-12 cycles of 98°C for 30s, 65°C for 30s, 72°C for 30s; and then 72 °C for 5min) with TransGen® Biotech FastPfu Fly DNA Polymerase and with VAHTS™ DNA Adapters for Illumina® (N302). PCR products were purified by GeneJET PCR Purification and eluted in 20-30ul EB. All the libraries were sequenced on Illumine X Ten platform.

### Quantitative PCR assays

Selected enhancer loci (Supplementary List 5) were PCR amplified from the genomic DNA of *A. thaliana*. Linearized vector pPBI221 was prepared by PCR amplification. Selected enhancer regions were cloned into the vector through Gibson Assembly reactions and verified by Sanger sequencing. Individual construct was tested in replicates by co-transfecting 1×10^6^ cells with the construct and an internal reference plasmid pGL3-SCP1-Luc-Basic. 24hrs post-transfection, total RNA was isolated from transfected cells with TransZol Up Plus kit (Transgen; cat. no. ER501-01) and reverse transcribed with TransScript II All-in-One first-strand cDNA Synthesis Supermix for qPCR (Transgen; cat. no. AH341). Using the QuantiNovaTM SYBR Green PCR kit (Qiagen; cat. no. 208052), we measured GFP relative expression at a CFX-96 Touch Real-Time PCR Detection System (Bio-Rad; cat. no. 1855195) in triplicates for each sample with the following program: 95°C 2min, followed by 40 cycles of 95°C 5s and 60°C 10s, 4°C hold.

### STARR-seq data processing and library quality analysis

For STARR-seq sequencing data, we used bowtie2^57^ to align the reads to the *Arabidopsis thaliana* genome (TAIR10) with parameters “--no-discordant -X 2000”. Duplicate reads were removed by picard MarkDuplicates. Low quality reads were filtered by samtools view^58^ with parameters “-f 2 –q 5”. Two replicates of cDNA and plasmid were merged separately for subsequent peak calling. The correlation between replicates was calculated by deeptools^59^. The fragment size and coverage of STARR-seq library were calculated by picard CollectInsertSizeMetrics and CollectWgsMetricsWithNonZeroCoverage. The merged bam files were converted into bigwig format for subsequent IGV visualization.

For iSTARR-seq sequencing data, cDNA libraries contain the reads with oligo dT (aSTs reads) and without oligo dT (dSTs reads). Because oligo dT sequences are located in 5’ end of pair-end read2. We cut the 20bp at 5’ end and 30bp low quality bases at 3’ end for both pair-end reads. Then we carried out the same mapping, filtering, merging as in STARR-seq data processing. iSTARR-seq plasmid library contains 6 sub-libraries marked by 6 different index in each replicate. Each index library’s reads were mapped and filtered separately. And then we merged these reads from 6 index libraries into one bam file. The correlation, fragment size and coverage were also calculated like in STARR-seq.

### Enhancer identification

We first used R package BasicSTARRseq^18^ to call enhancers with parameters “minQuantile = 0.9, peakWidth = 500, maxPval = 0.001”. We calculated FDR for each identified genomic regions identified by BasidSTARRseq with ranked P values. For these regions, we further calculated their enhancer activity in two ways: enhancer activity in the full genomic region = (cDNA reads from the identified genomic region/total cDNA reads)/(plasmid reads from the identified genomic region/total plasmid reads); enhancer activity at the summit = (normalized cDNA reads at summit in identified genomic region)/(normalized plasmid reads corresponding to region of the cDNA summit). For an identified genomic region, if its enhancer activity in the full genomic region is bigger than 1, its enhancer activity at the summit is bigger or equals to 1.3 and with a p vale smaller than 0.001 and FDR smaller than 0.001 as well, then it is considered as an enhancer in this work.

### Chromatin accessibility and ChIP-seq data processing

We collected published chromatin accessibility data and ChIP-seq data in which experimental materials and developmental stages are similar to the *A. thaliana* material used for STARR-seq and iSTARR-seq in this study. These raw data were downloaded from published study in GEO database, including DNase-seq^48^, ATAC-seq^44^, H3K4me1^46^, H3K27ac^46^, H3K9ac^44^, H3K36ac^45^, H3K56ac^41^, H3K36me3^43^, H3K4me3^40^, H3K27me3^46^, H3K9me2^39^, H2A^47^, H2A.Z^47^, RNAPII^42^. We performed trimming, mapping and filtering to these epigenetic data for subsequent analysis.

### Bisulfite-seq analysis

Bisulfite-seq raw data was downloaded from published study^60^ in GEO database. We used bismark^61^ to build TAIR10 index and map reads. The methylation levels were calculated by bismark_methylation_extractor with parameters “--paired-end --no_overlap --comprehensive --parallel 16 --bedGraph --buffer_size 20G --cytosine_report”.

### RNA-seq analysis

RNA-seq raw data was downloaded from published study^40^ in GEO database. We aligned reads to TAIR10 by hisat2^62^. The alignments were assembled into transcripts by Stringtie^63^. Gene expression was calculated by R package ballgown^64^.

### Enhancer genomic distribution analysis

We divided *Arabidopsis* genome into 9 functional regions, including TSS-200bp, TSS+/-50bp, 5’UTR, first intron, other introns, CDS, 3’UTR, intergenic and repeat regions. The ratio of every functional region relative to the whole genome was calculated. Enhancer distribution in genome was analyzed by overlapping enhancer midpoint with these functional regions. For enhancer distribution in TSS-genebody-TTS, gene proximal regions were divided into 3 parts: TSS upstream, gene body and TTS downstream. We divided each gene body into 40 equal bins to align genes of different lengths. TSS upstream and TTS downstream were extended by 50% of the average length of total genes and were divided into 20 equal bins. Then we calculated the coverage of enhancers in each bin of TSS-genebody-TTS by bedtools^65^. Finally, enhancer distribution in whole-genome was calculated by accumulating their distribution relative to each gene’s TSS-genebody-TTS. Simultaneously, we randomly selected the same number of sites from the whole genome as control group to calculate their distribution pattern relative to TSS-genebody-TTS.

### Enhancer with gene expression analysis

We assigned proximal genes to enhancers which are within 5kb from TSS to the midpoint in an enhancer. Genes can be classified into 2 types based on the existence of enhancer or not. Enhancer density in gene proximal 5kb regions was calculated. To reveal the relationship between gene expression and enhancer distribution, we classified genes into 4 categories by whether enhancers existed in their TSS region (from TSS-500bp to TSS+100bp) and TTS region (TTS+/-300bp) or not. We assigned the nearest gene (distance between TSS to enhancer midpoint) to each enhancer and compared the expression of genes. We used clusterProfiler^66^ to perform gene ontology (GO) analysis for genes expressed at different levels with enhancer in proximity. GO term enrichment was cut off by P value less than 0.05. And the results showed top 5 enriched GO term ranked by enrichment and P value in each gene expression level.

### Epigenetic signal enrichment at enhancers

Epigenetic signal enrichment at enhancers was computed by deeptools. We used deeptools computeMatrix to calculate epigenetic signal enrichment in each bin of enhancer-proximal 3kb regions with parameters “--referencePoint center -b 3000 -a 3000 –skipZeros --binSize 100 --averageTypeBins sum”. The heatmap was plotted by deeptools plotHeatmap in a descending order of average signal strength in each enhancer.

### Enhancer cluster on epigenetic signals

Enhancers were classified into 4 categories on their epigenetic signals in this study. First, we computed the coverage of epigenetic data’s aligned reads in each enhancer by bedtools to obtain the matrix of epigenetic enrichment signal at enhancers. We then performed k-means clustering for enhancers according to their normalized epigenetic signal strength. Finally, enhancer cluster count was confirmed according to the sum of squares within groups in different k-means cluster counts and enhancer actual epigenetic signals. Each enhancer cluster’s genomic distribution and proximal gene expression were also analyzed.

### Transcription factor binding sites (TFBSs) enrichment in enhancers

We obtained 529 *A. thaliana* transcription factor binding site data generated by DAP-seq in a previously published study^50^. All TFBSs were overlapped with enhancers to calculate their enriched counts by bedtools. Transcription factors belonging to a same big family were combined together. Total 43 datasets of transcription factors binding sites were kept after combining and filtering. We randomly selected genomic sites (same number as enhancers) as control group to perform the same computation. The relative enrichment of TFBSs was computed by comparing the number of them in enhancers and in random sites. The P value was calculated by Fisher’s exact test for each TFBS.

### Chromatin interaction analysis

Chromatin interaction data in *A. thaliana* were downloaded from previous publication by Liu et. al.^29^. We used chromatin loop anchors to overlap with enhancers and promoters. Based on these results, we analyzed the frequency of chromatin interactions between enhancer-enhancer, enhancer-promoter and enhancer-other regions.

### Statistical analysis

We used R for all statistical analysis.

## Data Availability

All raw data are deposited in the sequence read archive under the accession number GSE157030 and are accessible with secure token etqxquucrbcbxcj. All raw data are also available under the accession number of CRA003161 at http://bigd.big.ac.cn/gsa/s/cYaC5mpq.

## Code Availability

Custom codes are available at https://github.com/jing-wan/iSTARR-seq_data_analysis.

## Authors’ contributions

C.H. and L.N. conceived and designed the project. L.N. and J.S. constructed the reporter library and validated identified sites. J.W. and N.H. carried out bioinformatics analysis. Y.H. participated in cell preparation, transfection and sequencing library preparation. L.L. and C.H. supervised the data analysis. C.H. supervised the project. C.H. wrote the manuscript with input from all authors.

## Acknowledgements

We gratefully acknowledge financial support from the National Key R&D Program of China (2018YFC1004500), the National Natural Science Foundation of China (31571347 to CH and 31771430 to LL), Southern University of Science and Technology (No.G02226301 and Y01501821 to C.H.) and Huazhong Agricultural University Scientific & Technological Self-innovation Foundation (to LL), and support from the Center for Computational Science and Engineering of Southern University of Science and Technology.

## Figure legends

**Supplementary Table 1 | Sequencing statistics of STARR-seq and iSTARR-seq plasmid and cDNA libraries.**

**Supplementary Table 1:**
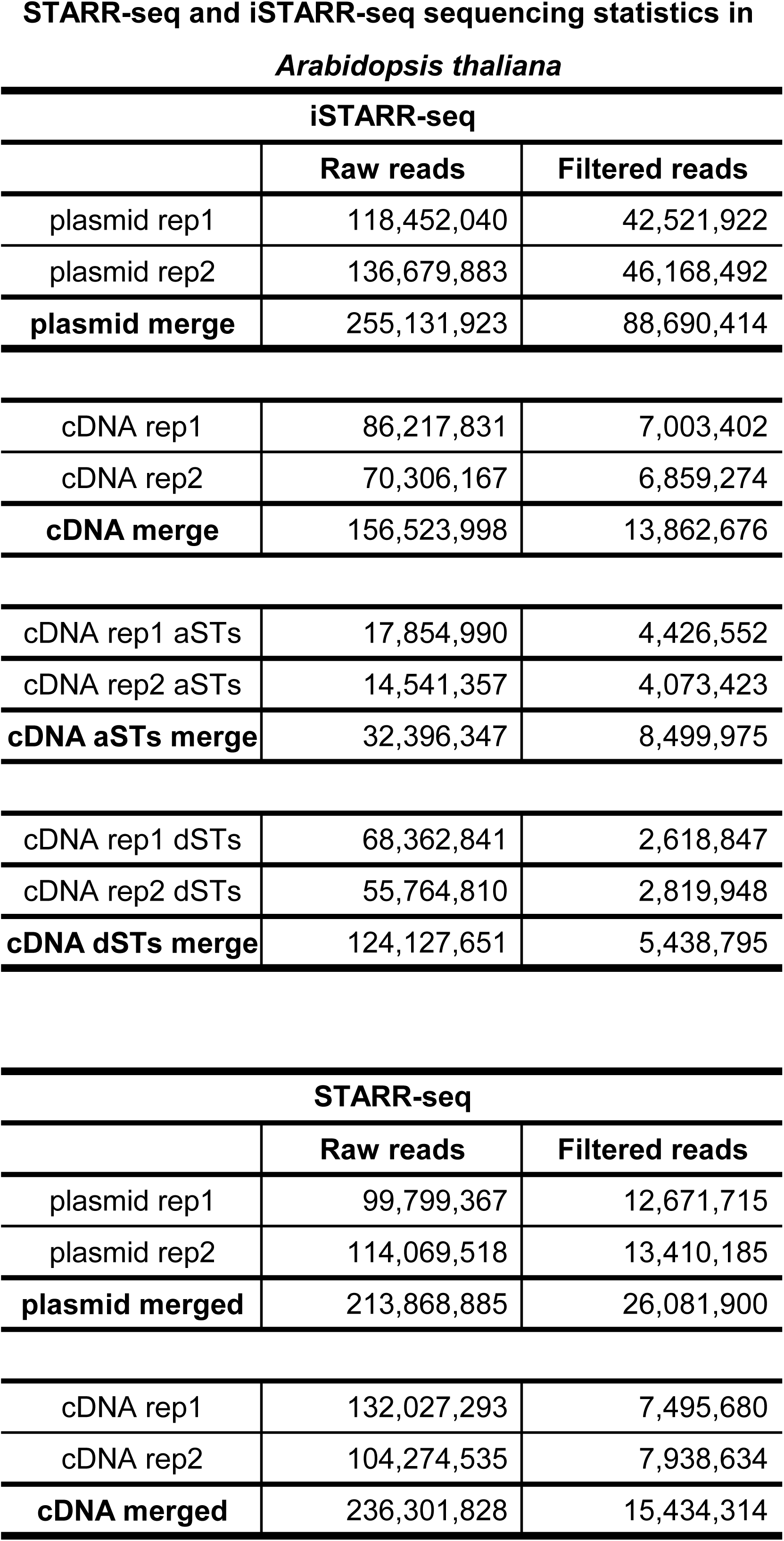
STARR-seq and iSTARR-seq sequencing statistics in *Arabidopsis thaliana*.

| iSTARR-seq |  |  |
| --- | --- | --- |
|  | Raw reads | Filtered reads |
| plasmid rep1 | 118,452,040 | 42,521,922 |
| plasmid rep2 | 136,679,883 | 46,168,492 |
| **plasmid merge** | **255,131,923** | **88,690,414** |
| cDNA rep1 | 86,217,831 | 7,003,402 |
| cDNA rep2 | 70,306,167 | 6,859,274 |
| **cDNA merge** | **156,523,998** | **13,862,676** |
| cDNA rep1 aSTs | 17,854,990 | 4,426,552 |
| cDNA rep2 aSTs | 14,541,357 | 4,073,423 |
| **cDNA aSTs merge** | **32,396,347** | **8,499,975** |
| cDNA rep1 dSTs | 68,362,841 | 2,618,847 |
| cDNA rep2 dSTs | 55,764,810 | 2,819,948 |
| **cDNA dSTs merge** | **124,127,651** | **5,438,795** |
| STARR-seq |  |  |
|  | Raw reads | Filtered reads |
| plasmid rep1 | 99,799,367 | 12,671,715 |
| plasmid rep2 | 114,069,518 | 13,410,185 |
| **plasmid merged** | **213,868,885** | **26,081,900** |
| cDNA rep1 | 132,027,293 | 7,495,680 |
| cDNA rep2 | 104,274,535 | 7,938,634 |
| **cDNA merged** | **236,301,828** | **15,434,314** |

**Supplementary Fig. 1.**
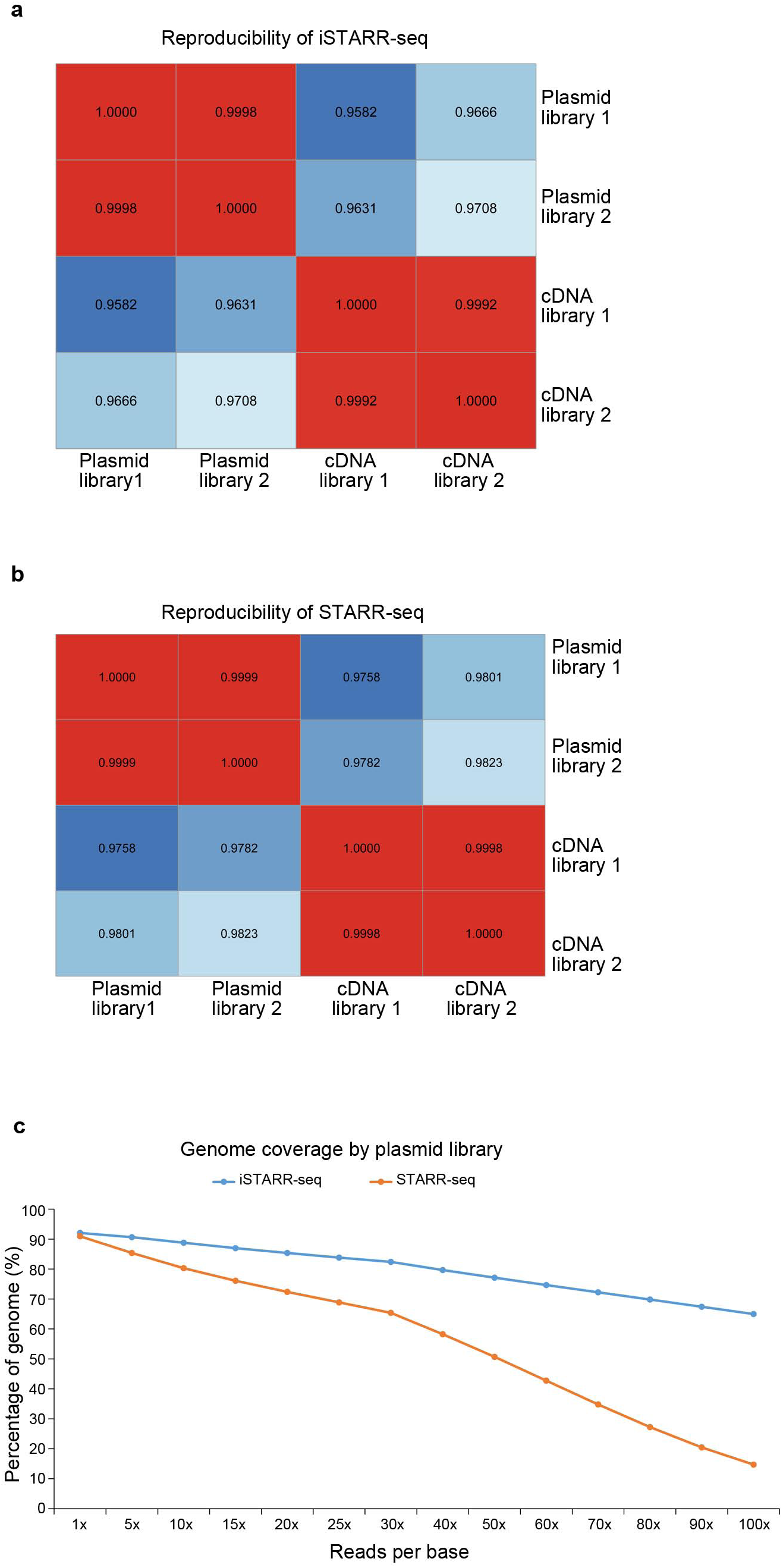
Reproducibility and quality of STARR-seq and iSTARR-seq libraries. Pearson’s correlation coefficients were calculated for all libraries of iSTARR-seq **(a)** and STARR-seq **(b)** experiments. **c**, The times that a single nucleotide being covered by reads from the plasmid libraries were calculated for the whole genome.

**Supplementary Fig. 2.**
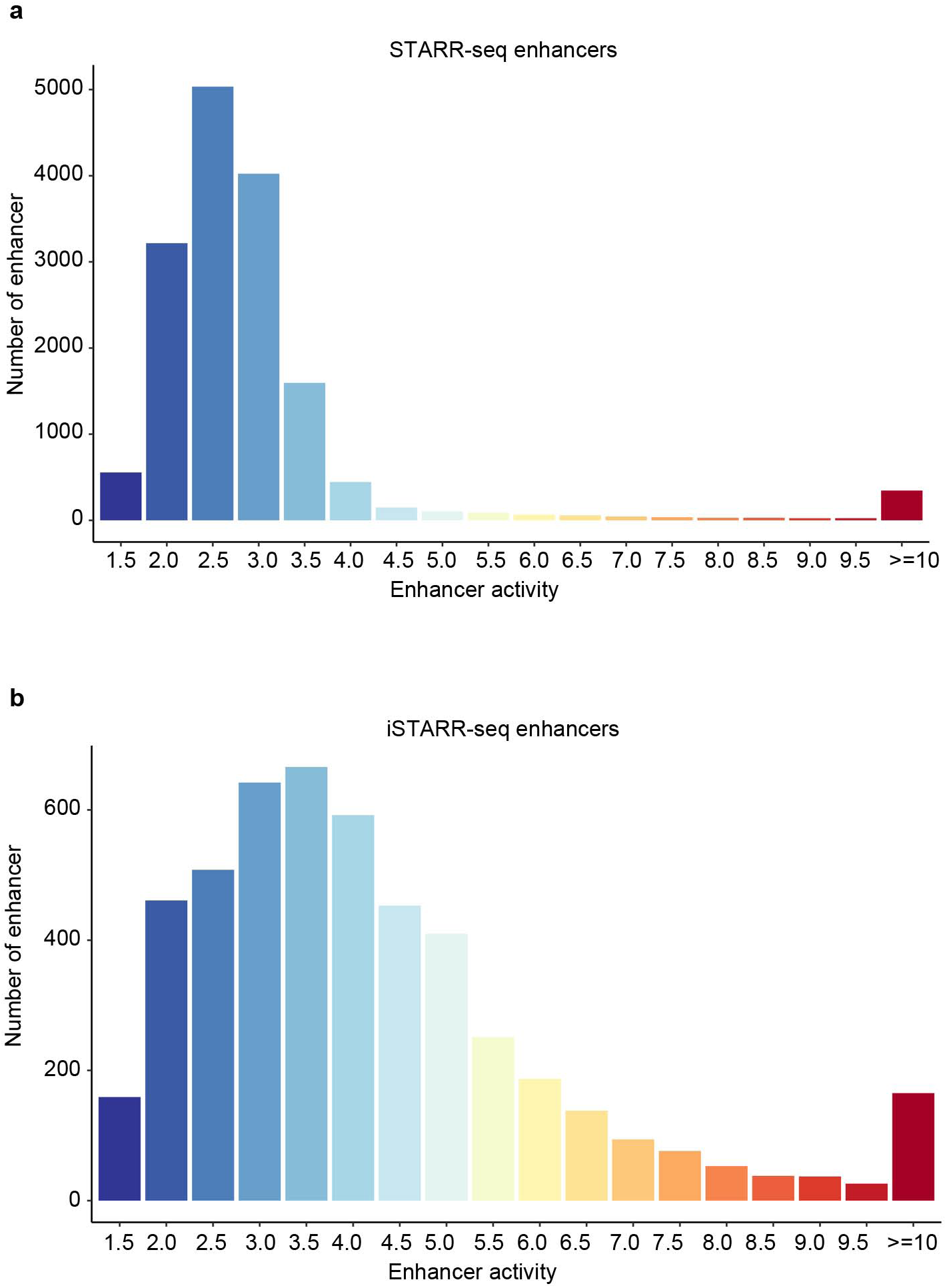
STARR-seq and iSTARR-seq enhancers. Distribution of enhancers activity for STARR-seq (**a**) and iSTARR-seq (**b**) experiments.

**Supplementary Fig. 3.**
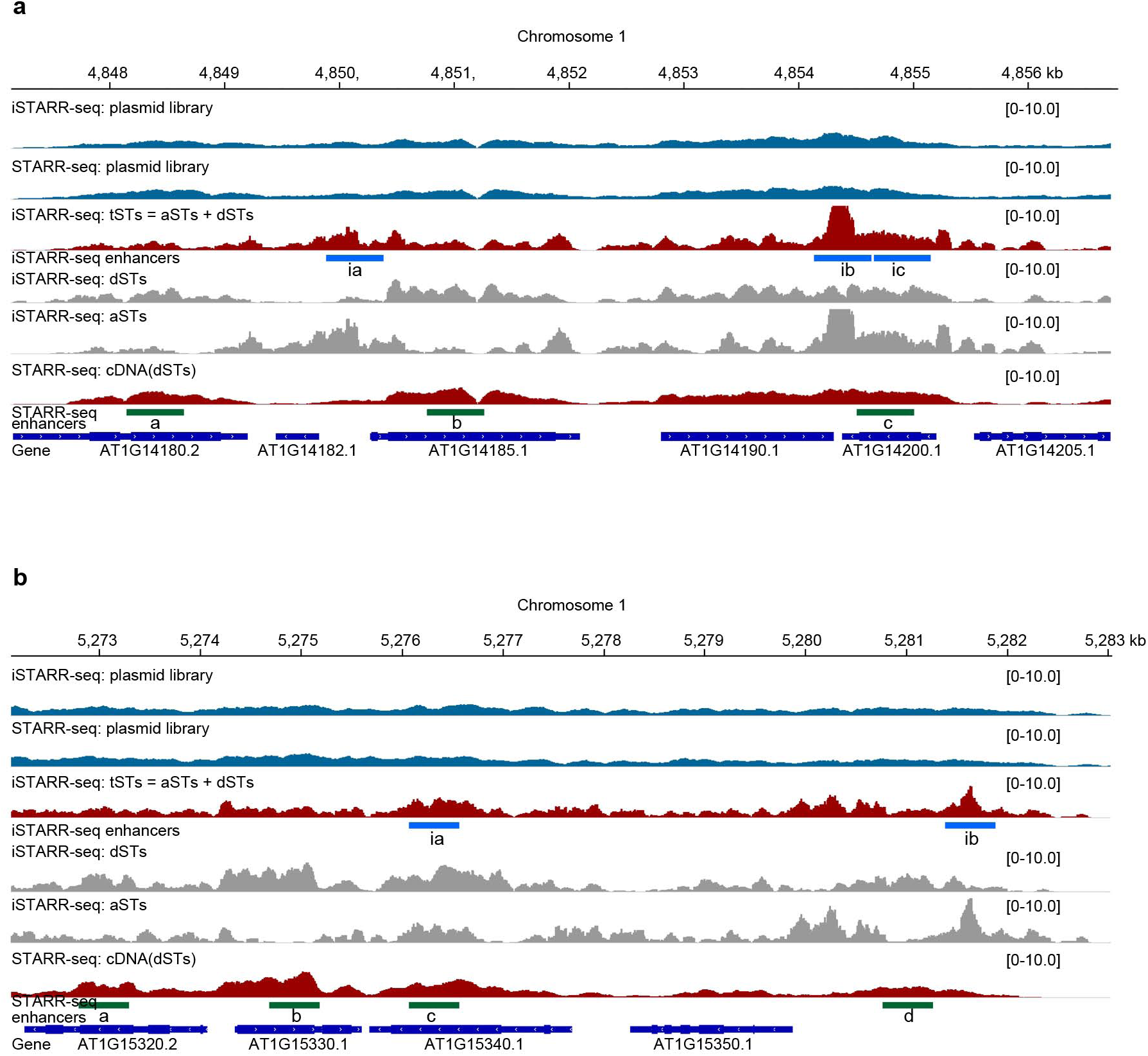
Two example regions showing STARR-seq and iSTARR-seq enhancers identified (a, b).

**Supplementary Fig. 4.**
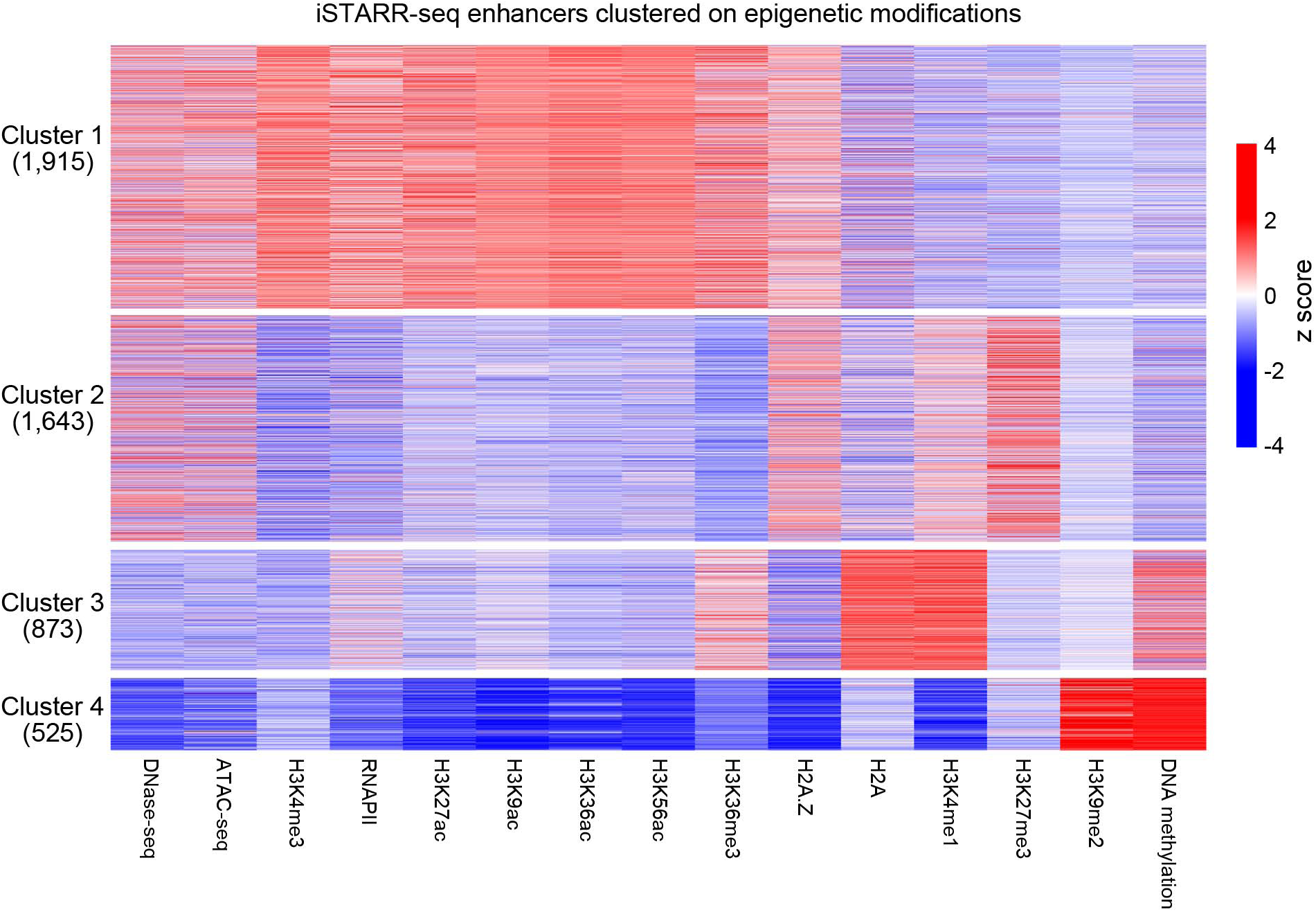
Clusters of iSTARR-seq enhancers. iSTARR-seq enhancers were clustered on the combinatory epigenetic states at each enhancer.

**Supplementary List 1**|**Regions containing APAS identified in *A. thaliana*.**

**Supplementary List 2**| **Enhancers identified by STARR-seq**.

**Supplementary List 3**| **Enhancers identified by iSTARR-seq**.

**Supplementary List 4**|**Epigenetic datasets**.

**Supplementary List 5**|**In vitro validated enhancers**.

## References

1. Maston, G.A., Landt, S.G., Snyder, M. & Green, M.R. Characterization of Enhancer Function from Genome-Wide Analyses. Annu Rev Genom Hum G 13, 29–57 (2012).

2. Ong, C.T. & Corces, V.G. Enhancer function: new insights into the regulation of tissue-specific gene expression. Nature Reviews Genetics 12, 283–293 (2011).

3. Bulger, M. & Groudine, M. Enhancers: The abundance and function of regulatory sequences beyond promoters. Developmental Biology 339, 250–257 (2010).

4. Tang, Z. et al. CTCF-Mediated Human 3D Genome Architecture Reveals Chromatin Topology for Transcription. Cell 163, 1611–1627 (2015).

5. Kieffer-Kwon, K.R. et al. Interactome maps of mouse gene regulatory domains reveal basic principles of transcriptional regulation. Cell 155, 1507–1520 (2013).

6. Li, G. et al. Extensive promoter-centered chromatin interactions provide a topological basis for transcription regulation. Cell 148, 84–98 (2012).

7. Bulger, M. & Groudine, M. Functional and mechanistic diversity of distal transcription enhancers. Cell 144, 327–339 (2011).

8. Whyte, W.A. et al. Master transcription factors and mediator establish super-enhancers at key cell identity genes. Cell 153, 307–319 (2013).

9. Visel, A. et al. A high-resolution enhancer atlas of the developing telencephalon. Cell 152, 895–908 (2013).

10. Hnisz, D. et al. Super-enhancers in the control of cell identity and disease. Cell 155, 934–947 (2013).

11. Zentner, G.E., Tesar, P.J. & Scacheri, P.C. Epigenetic signatures distinguish multiple classes of enhancers with distinct cellular functions. Genome Res 21, 1273–1283 (2011).

12. Kim, T.K. et al. Widespread transcription at neuronal activity-regulated enhancers. Nature 465, 182–187 (2010).

13. Creyghton, M.P. et al. Histone H3K27ac separates active from poised enhancers and predicts developmental state. Proc Natl Acad Sci U S A 107, 21931–21936 (2010).

14. Visel, A., Rubin, E.M. & Pennacchio, L.A. Genomic views of distant-acting enhancers. Nature 461, 199–205 (2009).

15. Visel, A. et al. ChIP-seq accurately predicts tissue-specific activity of enhancers. Nature 457, 854–858 (2009).

16. Heintzman, N.D. et al. Histone modifications at human enhancers reflect global cell-type-specific gene expression. Nature 459, 108–112 (2009).

17. Heintzman, N.D. et al. Distinct and predictive chromatin signatures of transcriptional promoters and enhancers in the human genome. Nat Genet 39, 311–318 (2007).

18. Arnold, C.D. et al. Genome-wide quantitative enhancer activity maps identified by STARR-seq. Science 339, 1074–1077 (2013).

19. Muerdter, F. et al. Resolving systematic errors in widely used enhancer activity assays in human cells. Nat Methods 15, 141–149 (2018).

20. Liu, Y. et al. Functional assessment of human enhancer activities using whole-genome STARR-sequencing. Genome Biol 18, 219 (2017).

21. Liu, S. et al. Systematic identification of regulatory variants associated with cancer risk. Genome Biol 18, 194 (2017).

22. Dao, L.T.M. et al. Genome-wide characterization of mammalian promoters with distal enhancer functions. Nat Genet 49, 1073–1081 (2017).

23. Vanhille, L. et al. High-throughput and quantitative assessment of enhancer activity in mammals by CapStarr-seq. Nat Commun 6, 6905 (2015).

24. Inoue, F. & Ahituv, N. Decoding enhancers using massively parallel reporter assays. Genomics 106, 159–164 (2015).

25. Sun, J. et al. Global Quantitative Mapping of Enhancers in Rice by STARR-seq. Genomics Proteomics Bioinformatics 17, 140–153 (2019).

26. Zhu, S. et al. PlantAPAdb: A Comprehensive Database for Alternative Polyadenylation Sites in Plants. Plant Physiology 182, 228–242 (2020).

27. Wu, X.H. et al. Genome-wide characterization of intergenic polyadenylation sites redefines gene spaces in Arabidopsis thaliana. Bmc Genomics 16 (2015).

28. Wu, X.H. et al. Genome-wide landscape of polyadenylation in Arabidopsis provides evidence for extensive alternative polyadenylation. P Natl Acad Sci USA 108, 12533–12538 (2011).

29. Liu, C. et al. Genome-wide analysis of chromatin packing in Arabidopsis thaliana at single-gene resolution. Genome Research 26, 1057–1068 (2016).

30. Hsieh, T.H.S. et al. Mapping Nucleosome Resolution Chromosome Folding in Yeast by Micro-C. Cell 162, 108–119 (2015).

31. Mukundan, B. & Ansari, A. Srb5/Med18-mediated Termination of Transcription Is Dependent on Gene Looping. Journal of Biological Chemistry 288, 11384–11394 (2013).

32. Tan-Wong, S.M. et al. Gene Loops Enhance Transcriptional Directionality. Science 338, 671–675 (2012).

33. Moabbi, A.M., Agarwal, N., El Kaderi, B. & Ansari, A. Role for gene looping in intron-mediated enhancement of transcription. P Natl Acad Sci USA 109, 8505–8510 (2012).

34. Tan-Wong, S.M., Wijayatilake, H.D. & Proudfoot, N.J. Gene loops function to maintain transcriptional memory through interaction with the nuclear pore complex. Gene Dev 23, 2610–2624 (2009).

35. Laine, J.P., Singh, B.N., Krishnamurthy, S. & Hampsey, M. A physiological role for gene loops in yeast. Gene Dev 23, 2604–2609 (2009).

36. Singh, B.N. & Hampsey, M. A transcription-independent role for TFIIB in gene looping. Molecular Cell 27, 806–816 (2007).

37. O’Sullivan, J.M. et al. Gene loops juxtapose promoters and terminators in yeast. Nature Genetics 36, 1014–1018 (2004).

38. Lee, J.Y. et al. Transcriptional and posttranscriptional regulation of transcription factor expression in Arabidopsis roots. P Natl Acad Sci USA 103, 6055–6060 (2006).

39. Zhao, S. et al. Plant HP1 protein ADCP1 links multivalent H3K9 methylation readout to heterochromatin formation. Cell Res 29, 54–66 (2019).

40. Zhang, H. et al. The effects of Arabidopsis genome duplication on the chromatin organization and transcriptional regulation. Nucleic Acids Research 47, 7857–7869 (2019).

41. Lu, Z.F. et al. The prevalence, evolution and chromatin signatures of plant regulatory elements. Nature Plants 5, 1250–1259 (2019).

42. Chen, C. et al. RNA polymerase II-independent recruitment of SPT6L at transcription start sites in Arabidopsis. Nucleic Acids Research 47, 6714–6725 (2019).

43. Wollmann, H. et al. The histone H3 variant H3.3 regulates gene body DNA methylation in Arabidopsis thaliana. Genome Biology 18 (2017).

44. Jegu, T. et al. The Arabidopsis SWI/SNF protein BAF60 mediates seedling growth control by modulating DNA accessibility. Genome Biology 18 (2017).

45. Mahrez, W. et al. H3K36ac Is an Evolutionary Conserved Plant Histone Modification That Marks Active Genes. Plant Physiology 170, 1566–1577 (2016).

46. Zhu, B., Zhang, W.L., Zhang, T., Liu, B. & Jiang, J.M. Genome-Wide Prediction and Validation of Intergenic Enhancers in Arabidopsis Using Open Chromatin Signatures. Plant Cell 27, 2415–2426 (2015).

47. Yelagandula, R. et al. The Histone Variant H2A. W Defines Heterochromatin and Promotes Chromatin Condensation in Arabidopsis. Cell 158, 98–109 (2014).

48. Zhang, W.L., Zhang, T., Wu, Y.F. & Jiang, J.M. Genome-Wide Identification of Regulatory DNA Elements and Protein-Binding Footprints Using Signatures of Open Chromatin in Arabidopsis. Plant Cell 24, 2719–2731 (2012).

49. Yan, W.H. et al. Dynamic control of enhancer activity drives stage-specific gene expression during flower morphogenesis. Nature Communications 10 (2019).

50. O’Malley, R.C. et al. Cistrome and Epicistrome Features Shape the Regulatory DNA Landscape. Cell 165, 1280–1292 (2016).

51. Snyder, M.P. et al. Perspectives on ENCODE. nature 583, 693–698 (2020).

52. Sethi, A. et al. Supervised enhancer prediction with epigenetic pattern recognition and targeted validation. Nature Methods 17, 807-+ (2020).

53. Niu, L. et al. Amplification-free library preparation with SAFE Hi-C uses ligation products for deep sequencing to improve traditional Hi-C analysis. Commun Biol 2, 267 (2019).

54. Jayavelu, N.D., Jajodia, A., Mishra, A. & Hawkins, R.D. Candidate silencer elements for the human and mouse genomes. nature communications 11 (2020).

55. Chen, P.Y., Wang, C.K., Soong, S.C. & To, K.Y. Complete sequence of the binary vector pBI121 and its application in cloning T-DNA insertion from transgenic plants. Mol Breeding 11, 287–293 (2003).

56. Yoo, S.D., Cho, Y.H. & Sheen, J. Arabidopsis mesophyll protoplasts: a versatile cell system for transient gene expression analysis. Nature Protocols 2, 1565–1572 (2007).

57. Langmead, B. & Salzberg, S.L. Fast gapped-read alignment with Bowtie 2. Nat Methods 9, 357–359 (2012).

58. Li, H. et al. The Sequence Alignment/Map format and SAMtools. Bioinformatics 25, 2078–2079 (2009).

59. Ramirez, F. et al. deepTools2: a next generation web server for deep-sequencing data analysis. Nucleic Acids Res 44, W160–165 (2016).

60. Yang, R. et al. The developmental regulator PKL is required to maintain correct DNA methylation patterns at RNA-directed DNA methylation loci. Genome Biology 18 (2017).

61. Krueger, F. & Andrews, S.R. Bismark: a flexible aligner and methylation caller for Bisulfite-Seq applications. Bioinformatics 27, 1571–1572 (2011).

62. Kim, D., Landmead, B. & Salzberg, S.L. HISAT: a fast spliced aligner with low memory requirements. Nature Methods 12, 357–U121 (2015).

63. Pertea, M. et al. StringTie enables improved reconstruction of a transcriptome from RNA-seq reads. Nat Biotechnol 33, 290-+ (2015).

64. Frazee, A.C. et al. Ballgown bridges the gap between transcriptome assembly and expression analysis. Nat Biotechnol 33, 243–246 (2015).

65. Quinlan, A.R. & Hall, I.M. BEDTools: a flexible suite of utilities for comparing genomic features. Bioinformatics 26, 841–842 (2010).

66. Yu, G.C., Wang, L.G., Han, Y.Y. & He, Q.Y. clusterProfiler: an R Package for Comparing Biological Themes Among Gene Clusters. Omics 16, 284–287 (2012).

